# MHC class II transactivator effects on local and systemic immune responses in an α-synuclein seeded rat model for Parkinson’s disease

**DOI:** 10.1101/2022.12.02.518821

**Authors:** Filip Fredlund, Itzia Jimenez-Ferrer, Kathleen Grabert, Lautaro Belfiori, Kelvin C. Luk, Maria Swanberg

**Affiliations:** Translational Neurogenetics Unit, Department of Experimental Medical Science, Lund University, Lund, Sweden; Inflammation and Stem Cell Therapy Group, Division of Clinical Neurophysiology, Department of Clinical Sciences, Lund University, Lund, Sweden; Toxicology Unit, Institute of Environmental Medicine, Karolinska Institute, Stockholm, Sweden; Center for Neurodegenerative Disease Research, Department of Pathology and Laboratory Medicine, University of Pennsylvania Perelman School of Medicine, Pennsylvania, PA, USA

**Keywords:** Parkinson’s disease, MHC class II transactivator protein, MHC Class II Genes, alpha-Synuclein, Tumor Necrosis Factor, Immune Responses, Neuroinflammation

## Abstract

**BACKGROUND:** Parkinson’s disease (PD) is characterized by alpha-synuclein (α-Syn) pathology, neurodegeneration and neuroinflammation. HLA variants associated with PD and α-Syn specific circulating CD4+ T lymphocytes in PD patients highlight the importance of antigen presentation in PD etiology. The class II transactivator (CIITA) is the major regulator of MHCII expression. Reduced *Ciita* levels significantly increase α-Syn pathology, nigrostriatal neurodegeneration and behavioral deficits in α-Syn seed-induced rat PD models.

**OBJECTIVE:** To characterize immune profiles associated with enhanced PD-like pathology observed in rats expressing lower *Ciita* levels (DA.VRA4) compared to the background (DA) strain.

**METHODS:** To model PD, we combined rAAV-mediated α-Syn overexpression in the substantia nigra with striatal injection of α-Syn pre-formed fibrils (PFF). Immune profiles in brain and blood were analyzed by flow cytometry and multiplexed ELISA in naïve rats, 4- and 8 weeks post rAAV injection.

**RESULTS:** Flow cytometry showed *Ciita*-dependent regulation of MHCII on microglia, brain macrophages and circulating myeloid cells. The MHCII-dependent microglial response peaked at 4 weeks post rAAV injection, whereas the MHCII levels in circulating myeloid cells peaked at 8 weeks. There was no major infiltration of macrophages or T lymphocytes into the CNS in response to α-Syn and only subtle *Ciita*- and/or α-Syn-dependent changes in the T lymphocyte compartment. Lower *Ciita* levels were consistently associated with higher TNF levels in serum.

**CONCLUSIONS:** These results suggest that *Ciita* regulates susceptibility to PD-like pathology through minor but detectable changes in resident and peripheral immune cells and TNF levels, and indicate that mild immunomodulatory therapies could have therapeutic effects in PD.

## INTRODUCTION

Parkinson’s disease (PD) is a progressive and incurable neurodegenerative disorder estimated to affect 2-3% of the population above the age of 65 [1]. Since the large majority of all PD cases have a multifactorial etiology, where genetics, lifestyle and environment are contributing factors, the disease is complex [2]. A characteristic feature of PD is the degeneration of dopaminergic neurons in the substantia nigra pars compacta (SN), intraneuronal inclusions containing alpha-synuclein (α-Syn), and neuroinflammation [3]. The neuroinflammatory process includes microglial activation, local upregulation of major histocompatibility complex II (MHCII), altered levels of pro-inflammatory cytokines in cerebrospinal fluid (CSF), as well as systemic changes in blood cytokine levels and lymphocyte populations [3]. Genetic association studies have identified single nucleotide polymorphisms in the human leukocyte antigen (*HLA*) locus that regulate the expression of MHCII to be associated with an increased risk of developing PD [4-6]. Recently, coding polymorphisms causing amino-acid changes in *HLA-D* haplotypes (*HLA-DRB1*4*) were also shown to be associated to PD with a protective effect [7]. Collectively, this indicates that both the quantity and quality of MHCII are involved in PD etiology. Since MHCII molecules present antigens to T lymphocytes and induce antigen-specific responses they serve as a link between the innate and adaptive immune systems [8].

A role of the adaptive immune system in PD etiology is supported by the presence of lymphocytes in post-mortem brain tissue from PD patients [9] and findings of α-Syn reactive CD4+ lymphocytes [10, 11] early in the disease process [12]. However, it is not clear if and how antigen presentation contributes to or protects from PD pathology. The level of MHCII on antigen-presenting cells is controlled by the class II transactivator (CIITA, also known as MHC2TA) and *in vivo* silencing of *Ciita* using shRNA has been shown to prevent neurodegeneration in a nigral α-Syn overexpression model of PD in mice [13]. In contrast, we have previously found that congenic rats with lower CIITA and MHCII levels due to naturally occurring variants in the *Ciita* gene promotor have more widespread α-Syn pathology, more nigrostriatal neurodegeneration, more activated microglia and enhanced motor deficits after nigral overexpression of α-Syn alone [14] or combined with striatal seeding with α-Syn pre-formed fibrils (PFF) [15]. Of note, genetic variants mediating lower *CIITA* gene and MHC-protein expression are also found in humans and are associated with increased susceptibility to multiple sclerosis, rheumatoid arthritis and myocardial infarction, further adding to the interest of studying *CIITA* in relation to PD [16].

The aim of this study was to investigate the effect of *Ciita* expression on peripheral and local immune responses during α-Syn seeded PD-like pathology. To do so, we used a recombinant adeno-associated viral vector (rAAV) nigral α-Syn overexpression rat model combined with striatal seeding of human PFF in two rat strains with different susceptibility to PD-like pathology due to different transcriptional activity of the *Ciita* gene. The congenic DA.VRA4 strain has lower transcription of *Ciita* and *Mhc2*-genes and increased susceptibility to PD-like pathology compared to the background strain, DA. Using a flow cytometric approach, we investigated both brain- and peripheral immune populations. We confirmed previous results that DA.VRA4 rats have lower MHCII expression in microglial cells compared to DA rats and we observe a peak microglia response at 4 weeks post rAAV injection in both strains. In addition to the local effects, we found lower MHCII levels on circulating myeloid cells, subtle changes in CD4+/CD8+ T-lymphocyte proportions in blood as well as higher levels of tumor necrosis factor (TNF) in serum in DA.VRA4 rats. Collectively, these results suggest that the levels of *Ciita* alter distinct immune populations and cytokine levels that in turn affect the susceptibility and severity of PD.

## MATERIALS AND METHODS

### Experimental design

To investigate the effects of differential expression of *Ciita* we used wt DA rats and a congenic DA.VRA4 rat strain with lower levels of *Ciita* and MHCII [14]. Male rats entered the study at 12±1 weeks of age and a total of 77 rats were included with 6-9 rats/group. We used a combination of viral overexpression of human α-Syn combined with seeding of human PFF, adapted from Thakur et al [17]. Rats were injected with a rAAV6 vector carrying human α-Syn [18] into the SN followed two weeks later by an injection of human α-Syn PFF in the striatum (Fig. 1) [15]. Empty a rAAV6 vector and vehicle (Dulbecco’s phosphate buffered saline, DPBS) was used as control. Animals were sacrificed at 4- and 8-weeks post nigral injection for collection of brain, blood (whole blood and serum) and CSF samples. 6 animals per strain and time point were used for flow cytometric analysis of brain and blood samples and 2-3 animals per strain and time point were used for qualitative IHC validation of α-Syn expression, tyrosine hydroxylase (TH) loss, α-Syn pathology and MHCII upregulation. Naïve rats (n=6 per strain) were sacrificed at 12±1 weeks of age. Two rats (one naïve DA and one DA.VRA4 α-Syn 8 week) were excluded from flow cytometry analysis of brain due to unsatisfactory perfusion and blood-filled ventricles, respectively. One DA rat from the 8-week α-Syn group was excluded from flow cytometry analysis for both blood and brain due to a clogged capillary during stereotactic surgery. One DA rat from the 8-week control group was excluded for flow cytometry analysis of blood due to inadequate number of events.

**Fig. 1.**
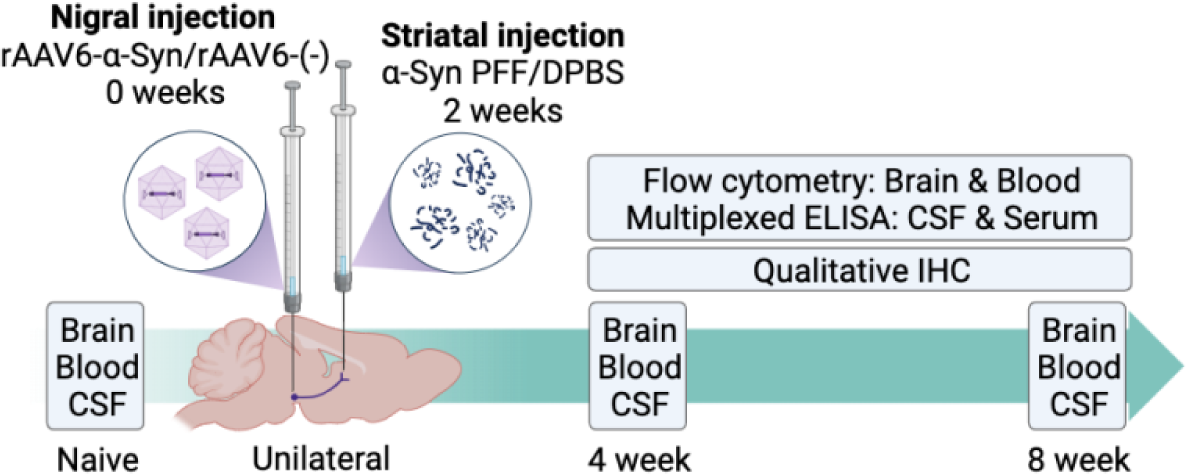
Experimental outline. Unilateral PD-like pathology was induced in DA and DA.VRA4 rats by rAAV6-mediated overexpression of human α-Syn in substantia nigra (week 0) combined with striatal seeding of human pre-formed fibrils of α-Syn (PFF, week 2). A control vector combined with vehicle was used as control (rAAV6-(-)+DPBS). Brain, blood and CSF were collected from naïve rats, at 4- and 8-weeks post nigral injection. Immune populations in brain and blood were characterized by flow cytometry and cytokine levels in serum and CSF were analyzed by multiplexed ELISA. Qualitative immunohistochemistry was performed to assess α-Syn pathology, as well as the level of neurodegeneration and microglial activation.

### Animals

The DA.VRA4 strain was generated by transfer of the VRA4 locus from the PVG strain to a DA background [19]. Rats were housed 2-3 per cage in “type III high” individually ventilated cages with free access to standard rodent chow and water and kept in a pathogen-free and climate-controlled environment with a 12 h light/dark cycle at the Biomedical Center in Lund. All procedures were approved by the local ethics committee in the Malmö-Lund region and specified in permit 18037-19.

### Viral vectors

rAAV6 carrying human α-Syn under transcriptional regulation by the Synapsin-1 promotor and the woodchuck hepatitis virus posttranscriptional regulatory element was generated as previously described [18] and injected at a concentration of 1.3E+10 gc/µl. The same vector but without the human α-Syn gene was used as a control and injected at a concentration of 1.7E+10 gc/µl. Concentration was determined by ITR-qPCR.

### Pre-formed fibrils

Human α-Syn PFF were produced as previously described[20] and stored at -80 °C until use. PFF were diluted to a concentration of 2.5 µg/µl in sterile DPBS and sonicated for 6 min with 1 s ON/1 s OFF pulses at 70% power using a Q125 sonicator and cup horn (Qsonica, U.S.). The gross structure of PFF before and after sonication were imaged using transmission electron microscopy. PFF were diluted to a concentration of 0.025 µg/µl and transferred to a hexagonal pattern 400 mesh cupper grid with a pioloform film reinforced with a carbon coat, for 20 min at room temperature (RT). Samples were stabilized with uranyl acetate for 1 min. Excess uranyl acetate was removed and the grids were left to dry for at least 5 min prior to imaging using a FEI Tecnai Spirit BioTWIN transmission electron microscope (FEI, U.S.).

### Surgical procedure

Rats were anaesthetized with 5% and maintained with 1-3% isoflurane (Isoflo vet, Orion Pharma) with a 2:1 mixture of O_2_: NO_2_ during the surgical procedure. Rats were attached to a stereotactic frame with a flat-skull position and 0.2 ml. Marcain (2.5 mg/ml, Aspen Nordic, Denmark) was subcutaneously (s.c.) injected under the scalp for local analgesia. Burr holes were created using a dental drill. For nigral injections, 3 µl rAAV6-(-) or rAAV6-α-Syn was injected in the following coordinates taken from bregma [21]; Anterior/posterior (A/P) -5.3 mm, medial/lateral (M/L) ±1.7 mm and dorsal/ventral (D/V) -7.2 mm. For striatal injections, 3 µl of sonicated PFF (2.5 µg/µl) or DPBS as control was injected using the following coordinates relative to bregma [21]; A/P -0.4 mm, M/L ±3.0 mm and D/V -4.5 mm. Injections were made unilaterally in the right hemisphere (ipsilateral) using a 10 µl Hamilton syringe (Hamilton, U.S.) fitted with a glass capillary. Injections were made with a flow rate of 0.5 µl/2 min and the capillary was left for 2 min after the injection before it was slowly retracted. The wound was sutured using surgical staples. Metacam (1 mg/kg) (Boehringer Ingelheim Animal Health, Germany) was injected s.c. for post-operative analgesia. The rats were left to recover in clean cages and monitored for 48 h post-surgery.

### Tissue collection

Rats were euthanized by intraperitoneal injection of 200-300 mg/kg sodium pentobarbital (APL, Sweden). Samples were collected in the following order: CSF, blood (serum and whole blood) and brain. CSF and blood were collected prior to perfusion whereas brain samples were collected after perfusion.

#### Cerebrospinal fluid (CSF) sampling

CSF samples were collected at baseline, 4- and 8-weeks post nigral injection from all 77 rats in a stereotactic frame with an approximate 50-60° downward flex of the head. A midline incision was made over the neck and muscles covering the cisterna magna were severed using a scalpel. CSF samples were aspirated using a 27G scalp vein set (Vygon, France) by inserting the bevel of the needle perpendicular to the cisterna magna. CSF was collected into protein LoBind tubes (Eppendorf, Germany), immediately put on dry ice and stored at -80 °C until analysis. CSF samples contaminated with blood were excluded from analysis.

#### Serum and whole blood collection

Blood from naïve, 4- and 8-week time points from all 77 rats included in the study was collected by cardiac puncture. For cytokine analysis, serum was prepared by leaving whole blood undisturbed at RT for 30-60 min followed by centrifugation for 10 min at 4 °C and 2,000xg. Serum was aliquoted into protein LoBind tubes (Eppendorf, Germany) and stored at -80 °C until analysis. Whole blood was collected into K3E EDTA coated tubes (BD, U.S.) and stored at 4 °C for 3-4 h until preparation for flow cytometric analysis.

#### Brain processing for immunohistochemistry and flow cytometry

After CSF and blood sampling, rats were transcardially perfused with 0.9% saline (w/v) with the descending aorta clamped using hemostatic forceps for at least 5 min or until no blood was visible. For IHC analysis, rats were subsequently perfused with ice-cold 4% paraformaldehyde (PFA) (w/v) for 5 min and the brains post-fixed in 4% PFA (w/v) at 4 °C overnight followed by cryopreservation in PBS containing 30% sucrose (w/v) and 0.01% sodium azide (w/v), pH 7.2 until sectioning. For flow cytometric analysis, saline-perfused brains were collected into ice-cold Roswell Park Memorial Institute 1640 medium without phenol red (Gibco/Thermo Fischer Scientific, U.S.) and stored at 4 °C for a maximum of 3 h until processing.

### Sample preparation for Flow cytometry

#### Brain sample collection and homogenization

Hemispheres of freshly collected brains were separated and put into a 7 ml glass dounce tissue grinder (DWK, Germany) with 3-5 ml ice-cold 1x Hank’s Balanced Salt Solution without calcium, magnesium or phenol red (HBSS) (Gibco/Thermo Fischer Scientific, U.S.), pH 7.0-7.4. Each hemisphere was homogenized on ice using the large clearance pestle followed by the small clearance pestle until complete homogenization. The glass dounce tissue grinder set was washed with detergent, rinsed and dried between samples. Homogenized samples were passed through a 100 µm nylon cell strainer (Falcon, U.S.) into a 50 ml conical tube to remove any remaining large debris. 1x HBSS (pH 7.0-7.4) was added until a total volume of 12 ml was reached and samples were kept on ice until separation of myelin and brain mononuclear cells.

#### Brain mononuclear cell isolation by gradient separation

Brain mononuclear cells were isolated and myelin removed using an adapted two-layer density gradient protocol [22, 23]. A 100% stock isotonic Percoll (SIP) was prepared by diluting Percoll (GE Healthcare, U.S.) 9:1 in 10x HBSS (Gibco/Thermo Fischer Scientific, U.S.) and 35% SIP was prepared by diluting 100% SIP 0.35:1 in 1x HBSS pH 7.0-7.4. Homogenized brain samples were centrifuged for 5 min at 4 °C and 400xg, the supernatant was discarded and the pellet was thoroughly resuspended in 16 ml of 35% SIP. The cell suspension was carefully layered with 5 ml of 1x HBSS pH 7.0-7.4 and centrifuged for 30 min at 4 °C and 800xg without brake. The HBSS layer (top), myelin layer (between HBSS and 35% SIP) and 35% SIP was aspirated and the pelleted isolated brain mononuclear cells were washed in 10 ml of 1x HBSS pH 7.0-7.4 and resuspended in ice-cold fluorescence-activated cell sorting (FACS) buffer.

#### Blood sample preparation

Whole blood samples collected in EDTA coated tubes was used for flow cytometric analysis (200 µl). Red blood cells were lysed by adding 1.8 ml of 1x Pharm Lyse (BD, U.S.) to whole blood cell samples and incubated at RT for 15-20 min. Cells were washed in sterile-filtered PBS (pH 7.2) and resuspended in sterile-filtered ice-cold FACS buffer (2% (w/v) bovine serum albumin fraction V (Roche, Switzerland) and 0.01% sodium azide (w/v) in PBS (pH 7.2)).

#### Antibody staining for flow cytometric analysis

FcγII receptors on blood and brain samples were blocked by adding anti-rat CD32 diluted 1:200 and incubated for 5 min at 4 °C. 50 µl of cell suspension was stained using an antibody cocktail (Table 1) diluted in Brilliant Stain Buffer (BD, U.S.). Cells were incubated with antibodies for 30 min at 4 °C in dark followed by washing in sterile PBS (pH 7.2). Cells were resuspended in 250 µl of sterile FACS buffer containing DRAQ7 diluted 1:1,000 prior to analysis.

**Table 1.** Antibodies, viability marker and compensation beads used for flow cytometry.

| Antigen/<br>Target | Species<br>specificity | Fluorochrom<br>e/<br>Conjugation | Clone | Isotype<br>/<br>Host | Dilutio<br>n | Company |
| --- | --- | --- | --- | --- | --- | --- |
| CD45 | Rat | APC-eFluor<br>780 | OX1 | Mouse<br>IgG1, κ | 1:100 | Invitrogen<br>(47-0461-82) |
| CD3 | Rat | BV421 | 1F4 | Mouse<br>IgM, κ | 1:200 | BD Horizon<br>(563948) |
| CD4 | Rat | BV605 | OX-35 | Mouse<br>IgG2a,<br>κ | 1:200 | BD OptiBuild<br>(740369) |
| CD8a | Rat | PE-Cy7 | OX8 | Mouse<br>IgG1, κ | 1:200 | Invitrogen<br>(25-0084-82) |
| CD11b | Rat | PE | WT.5 | Mouse<br>IgA, κ | 1:200 | BD<br>Pharmingen<br>(562105) |
| MHCII RT1B | Rat | Alexa Fluor<br>647 | OX-6 | Mouse<br>IgG1, κ | 1:400 | Bio-Rad<br>(MCA46A64<br>7) |
| CD86 | Rat | BV711 | 24F | Mouse<br>IgG1, κ | 1:100 | BD OptiBuild<br>(743215) |
| FcγRII | Rat | - | D34-485 | Mouse<br>IgG1, κ | 1:200 | BD<br>Pharmingen<br>(550270) |
| Compensati<br>on | Mouse, κ | - | - | - | - | BD<br>CompBeads<br>(552843) |
| Viability/<br>dsDNA | - | DRAQ7 | - | - | 1:1,000 | Invitrogen<br>(D15106) |

Samples were analyzed using an LSR Fortessa (BD, U.S.), configuration specified in Table 2. Compensation was performed using BD CompBeads (BD, U.S.) and prepared according to manufacturer’s instructions. Fluorescence minus one, unstained cells and unstained cells with viability dye were included for each recording session and for each sample type (blood or brain) and used to set gates. Gating strategy for brain and blood samples can be seen in Supplementary Fig. 2a and 3a. Microglial cells were gated as CD45^dim^CD11b+ in brain samples. Infiltrating macrophages/monocytes (CD45^high^CD11b+) and T lymphocytes (CD45+CD3+) in brain samples were rare with <1,000 events/hemisphere. Myeloid population in blood was gated as CD45+CD11b+ and T lymphocytes as CD45+CD3+. T helper cells were gated as CD4+ and cytotoxic T lymphocytes as CD8+. Data was analyzed using FlowJo software version 10.8.1 (BD, U.S.). All analyses were done on freshly isolated tissue and recorded during multiple sessions. 4-6 rats were used at each recording session (equal number of DA and DA.VRA4 rats per session) from the same experimental group (naïve/control/α-Syn) and time point (4- or 8-weeks).

**Table 2.** Configuration of the LSR Fortessa used for flow cytometric analysis and filters used for recording of isolated blood and brain cells.

| **Laser** | **Filter** | **Fluorochrome** |
| --- | --- | --- |
| Blue – 488 nm | 780/60 | PE-Cy7 |
|  | 695/40 | - |
|  | 610/20 | - |
|  | 575/26 | PE |
|  | 530/30 | - |
|  | 488/10 | SSC |
| Red – 640 nm | 780/60 | APC-eFluor 780 |
|  | 730/45 | DRAQ7 |
|  | 670/30 | Alexa Fluor 647 |
| Violet – 405 nm | 780/60 | - |
|  | 710/50 | BV711 |
|  | 660/20 | - |
|  | 610/20 | BV605 |
|  | 525/50 | - |
|  | 442/46 | BV421 |

### Immunohistochemistry

Fixed brains were coronally sectioned on a Microm HM450 freezing microtome (Thermo Scientific, U.S.) with 35 µm thickness in series of 12 and stored in Walter’s antifreeze solution at 4 °C until IHC staining. All stainings were done on free floating sections except for proteinase K treated human α-Syn staining which was done on sections mounted on gelatin-coated glass slides. Sections were rinsed with PBS or 0.1% PBS with Triton-X 100 (v/v) (PBST) between all incubation steps. For proteinase K resistant α-Syn aggregates, sections were incubated with 5 µg/ml Proteinase K (Thermo Fischer Scientific, U.S.) diluted in TBS for 1 h at RT prior to quenching. For 3,3’-diaminobenzidine (DAB) staining sections were quenched with 3% H_2_O_2_ (v/v) and 10% MetOH (v/v) in PBS. Sections were blocked with 10% serum (same species as secondary antibody) in 0.3% PBST. Primary antibody was diluted in 0.3% PBST with 5% serum (same species as secondary antibody) and incubated at 4 °C overnight. On the following day sections were incubated with biotinylated secondary antibody and incubated for 1 or 2 h at RT (DAB or Fluorescence, respectively). All antibodies used for IHC are found in Table 3. For DAB staining, horseradish peroxidase conjugated avidin/biotin-complex (Vector laboratories, U.S.) was prepared according to manufacturer’s instructions and added to the sections for 30 min at RT. A DAB substrate kit (Vector laboratories, U.S.) was prepared according to manufacturer’s instructions and used as a chromogen for visualization. DAB sections were mounted on gelatin-coated glass slides, dehydrated and coverslipped using Pertex (Histolab, Sweden). Fluorescently stained sections were coverslipped using PVA/DABCO and stored at 4 °C in dark. Brightfield overview images of TH and human α-Syn were acquired using an Olympus VS-120 virtual slide scanner (Olympus, Japan). Brightfield images of pS129 α-Syn and proteinase K treated human α-Syn in SN was acquired using an Olympus BX53 (Olympus, Japan). MHCII+ microglia cells were imaged using a Leica SP8 scanning confocal microscope (Leica, Germany).

**Table 3.** List of antibodies used for immunohistochemistry.

| **Antigen/Secondary antibody** | **Host** | **Dilution** | **Company** |
| --- | --- | --- | --- |
| Human α-Syn | Mouse | 1:1,000 | Santa Cruz (sc-12767) |
| Biotinylated anti-mouse | Horse | 1:200 | Vector Laboratories (BA-2001) |
| TH | Rabbit | 1:1,000 | EMD Millipore (AB152) |
| pS129 α-Syn | Rabbit | 1:2,000 | Abcam (ab51253) |
| Biotinylated anti-rabbit | Goat | 1:200 | Vector Laboratories (BA-1000) |
| MHCII | Mouse | 1:500 | Abcam (ab23990) |
| Alexa Fluor 488 anti-mouse | Donkey | 1:200 | Abcam (ab150105) |

### Cytokine analysis

Cytokine analysis in serum and CSF was performed using the V-PLEX Proinflammatory panel 2 Rat Kit from Mesoscale diagnostics (MSD, U.S.) according to manufacturer’s instructions. The plates were washed using PBS with 0.05% Tween-20 between incubation steps. Serum samples were diluted 4-fold and CSF samples 2-fold. Plates were read on a MESO QuickPlex SQ 120 analyzer (MSD, U.S.). Results were analyzed using the Discovery Workbench software version 4.0.13 (MSD, U.S.). The number of samples used for cytokine analysis differs as a consequence of available wells on the MSD plate. All samples were run in duplicates and the mean value was used for analysis. If only one replicate was detected it was included in the analysis. If both replicates were below the lower limit of detection (LLOD) for a sample the non-detected (ND) value was replaced with the lowest quantifiable value for the specific cytokine. If duplicates for more than one sample was below the LLOD for a group, no statistical comparisons were made and presented as non-detected (ND).

### Statistical analyses

Statistical analyses were conducted using the GraphPad Prism software version 10.1.1 (San Diego, CA, U.S.). Quantile-quantile plot of residuals was used to determine the use of parametric or non-parametric tests. Data in figures is presented as mean±SD and individual values. Naïve rats were compared by unpaired Student’s t-test. Groups at 4- and 8-weeks were compared with two-way ANOVA with Šídák multiple comparison test (DA vs. DA.VRA4 and control vs. α-Syn). Data in text is presented as (mean1±SD1 vs mean2±SD2, p-value, 95% CI of difference [lower limit, upper limit]). A significance level of α<0.05 was used for all analyses.

## RESULTS

### Qualitative immunohistochemistry confirms PD-like features characterized by α-Syn pathology and nigrostriatal dopaminergic neurodegeneration in the rAAV-α-Syn+PFF model

To investigate the effects of differential levels of *Ciita* on PD like-pathology we used the congenic DA.VRA4 rat strain with lower levels of *Ciita* and reduced expression MHCII [14] with DA rats as controls. Rats were injected with a rAAV-α-Syn vector [18] into the SN followed by an injection of sonicated human α-Syn PFF two weeks later in the striatum (α-Syn group) (Fig. 1 and Supplementary Fig. 1a). Control animals were injected with an empty rAAV vector into the SN and vehicle (DPBS) into the striatum (control group). Rats were sacrificed at baseline (naïve), 4- and 8-weeks post nigral injection for collection of brain, blood and CSF samples (Fig. 1).

The rAAV-α-Syn+PFF model used has been thoroughly characterized in a previous study, showing significantly reduced striatal TH+ fiber density and motor deficits in DA.VRA4 but not DA rats, as well as more aggregation and spread of α-Syn in DA.VRA4 compared to DA rats [15]. Quantification of the same parameters are not ethically warranted and not within the scope of the current study. Therefore, qualitative histological assessment was done to confirm the model. Robust staining for of human α-Syn was observed at 4- and 8-weeks in α-Syn (Supplementary Fig. 1b-c) but not in control groups (Supplementary Fig. 1d-e). As expected, the unilateral rAAV-α-Syn+PFF model induced loss of TH+ signal in the ipsilateral SN and striatum of both DA and DA.VRA4 rats at 4- and 8-weeks (Supplementary Fig. 1f-g), whereas the TH+ signal remained intact in the control groups (Supplementary Fig. 1h-i). rAAV-α-Syn+PFF injection also resulted in pathological forms of α-Syn aggregates, represented by positive signal for α-Syn phosphorylated at serine residue 129 (pS129) in cell somas and neurites, as well as by proteinase K-resistant α-Syn aggregates mainly observed as puncta along neurites in ipsilateral, but not contralateral SN (Supplementary Fig. 1j-k). Additionally, rAAV-α-Syn+PFF injection lead to upregulation of MHCII molecules in the ipsilateral but not contralateral midbrain of both DA and DA.VRA4 rats (Supplementary Fig. 1l).

### MHCII+ microglia peak at 4 weeks after rAAV-α-Syn+PFF and MHCII expression on microglia and brain macrophages is regulated by α-Syn and *Ciita*

Since DA.VRA4 rats with lower *Ciita* levels are more susceptible to α-Syn pathology and dopaminergic neurodegeneration than DA rats [14, 15], we characterized microglia, infiltrating macrophages and T lymphocyte populations in brain tissue by flow cytometry (Supplementary Fig. 2a). DA.VRA4 and DA did not differ in terms of microglial population size in the ipsilateral hemisphere except for naïve DA.VRA4 rats which had a lower percentage of microglia compared to DA in the right hemisphere (93±1.9% vs 96±1.1%, p=0.017, 95% CI [- 5.1, -0.65]) (Fig. 2a, b), however this difference was not observed in the left hemisphere (data not shown). In DA.VRA4, a lower amount of microglia was observed in the α-Syn group compared to controls at 4 weeks (93±1.4% vs 95±0.6%, p=0.042, 95% CI [-3.2, -0.06]) (Fig. 2b).

**Fig. 2.**
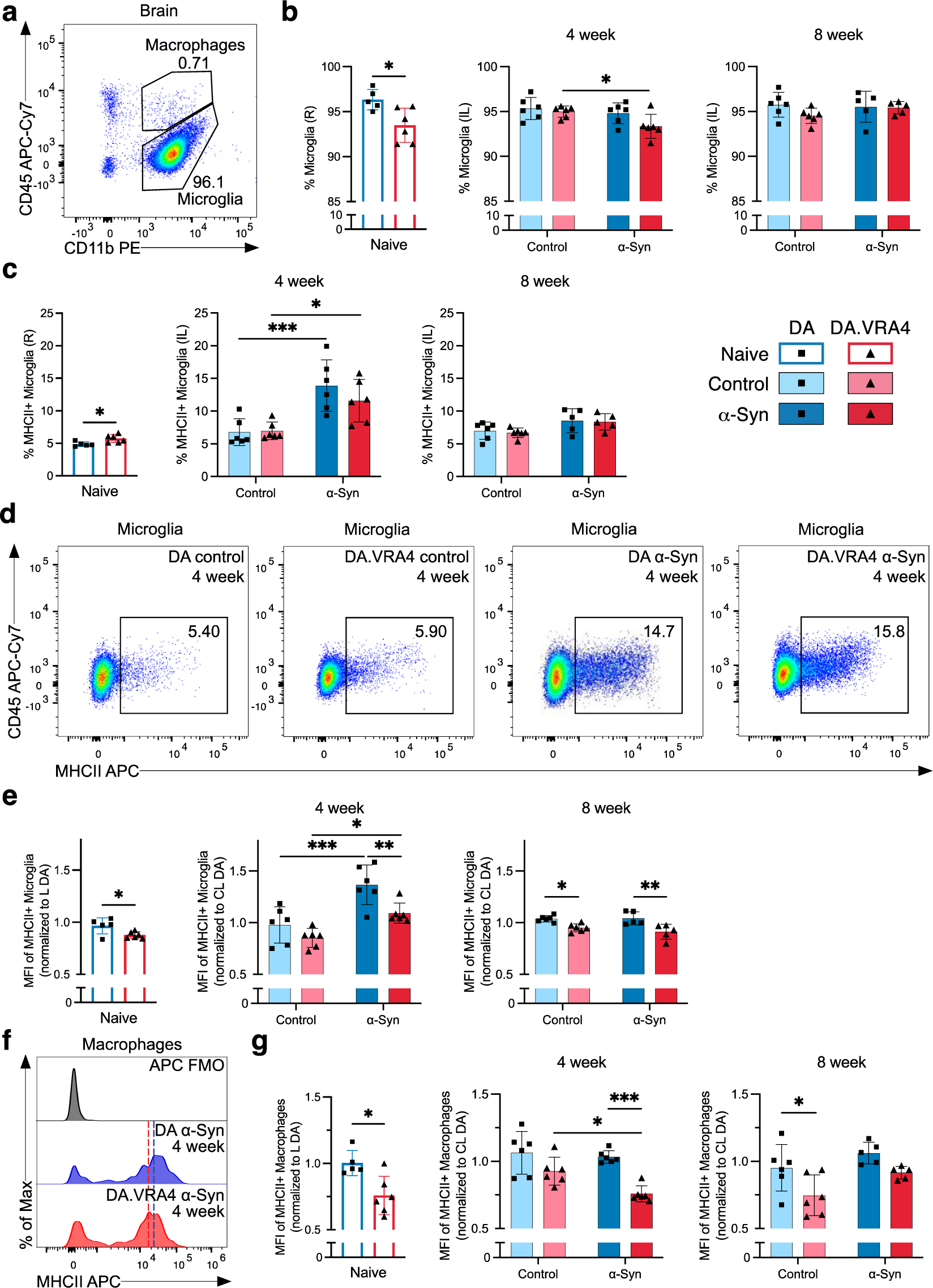
Effects of rAAV-α-Syn+PFF on microglia, brain infiltrating macrophages and MHCII expression. **a.** Gating of microglia (CD45^dim^CD11b+) and infiltrating macrophages/monocytes (CD45^high^CD11b+) in brain hemispheres. **b.** Quantification of microglia in right (R)/ipsilateral (IL) hemispheres. **c.** Quantification of MHCII+ microglia in R/IL hemispheres. **d.** Representative flow cytometric dot plots of MHCII+ microglia in IL hemispheres at the 4-week time point. **e.** Quantification of relative median fluorescence intensity (MFI) of MHCII+ microglia in R/IL hemispheres. At each recording session, R/IL MFI-values were normalized to the mean MFI-values in left (L)/contralateral (CL) hemisphere of DA. **f.** Representative histograms of MHCII expression on brain infiltrating macrophages with fluorescence minus one (FMO) control. The approximate MHCII+ MFI-value is indicated by dashed line; blue (DA) and red (DA.VRA4). **g.** Quantification of relative MFI of MHCII+ macrophages in R/IL hemispheres. As for microglia, R/IL MFI-values were normalized to the mean MFI-values of MHCII+ macrophages in L/CL hemisphere of DA for each recording session. **b, c, e and g.** Naïve (DA n=5, DA.VRA4 n=6), 4-week; control (DA n=6, DA.VRA4 n=6) and α-Syn (DA n=6, DA.VRA4 n=6), 8-week; control (DA n=6, DA.VRA4 n=6) and α-Syn (DA n=5, DA.VRA4 n=5). α-Syn=rAAV-α-Syn+PFF. Control=rAAV-(-)+DPBS. Naïve rats were compared by unpaired Student’s t-test. Groups at 4- and 8-weeks were compared with two-way ANOVA with Šídák multiple comparison test (DA vs. DA.VRA4 and control vs. α-Syn). * p < 0.05, ** p < 0.01 and *** p < 0.001. Data presented as mean ± SD with individual values.

In naïve DA.VRA4 rats, a larger proportion of microglia was MHCII+ compared to DA (5.7±0.60% vs 4.9±0.35%, p=0.019, 95% CI [0.18, 1.6]) (Fig. 2c). Although immunohistochemical staining has previously found increased numbers of activated microglia in the striatum of DA.VRA4 rats compared to DA [14, 15] the percentage of MHCII+ microglia in entire ipsilateral hemispheres did not differ between the strains at either 4-or 8-weeks (Fig. 2c). Compared to controls, rAAV-α-Syn+PFF injection lead to an expansion of MHCII+ microglia in both DA (14±4.0% vs 6.8±2.0%, p<0.001, 95% CI [3.2, 11]) and DA.VRA4 (12±3.2% vs 7.0±1.3%, p=0.021, 95% CI [0.65, 8.5]) at 4 weeks (Fig. 2c, d). By 8 weeks, the amount of MHCII+ microglia were reduced to baseline in both strains (Fig. 2c).

Quantification of MHCII on microglia (relative MFI-values; normalized to mean MHCII+ MFI in left/contralateral hemispheres of DA at each recording session) revealed that congenic DA.VRA4 rats had lower MHCII expression compared to DA in naïve rats (0.88±0.032 vs 0.97±0.077, p=0.029, 95% CI [-0.17, -0.011]), at 4 weeks in the α-Syn group (1.1±0.097 vs 1.4±0.19, p=0.008, 95% CI [-0.48, -0.070]) and at 8 weeks in both the control-(0.95±0.038 vs 1.0±0.030, p=0.019, 95% CI [-0.16, -0.014]) and α-Syn group (0.91±0.072 vs 1.0±0.063, p=0.002, 95% CI [-0.21, -0.05]) (Fig. 2e). Both strains had higher MHCII expression at 4 weeks in the α-Syn group compared to control (DA: 1.4±0.19 vs 0.98±0.18, p<0.001, 95% CI [0.19, 0.60] and DA.VRA4: 1.1±0.10 vs 0.85±0.09, p=0.020, 95% CI [0.04, 0.44]) (Fig. 2e). The increase in MHCII levels observed in α-Syn group at 4 weeks returned to control levels again at the 8-week time point in both strains (Fig. 2e).

We did not observe any strain- or α-Syn-dependent changes in infiltrating macrophages/monocytes (CD45^high^CD11b+) populations in brain in terms of overall amount or percentage of MHCII+ macrophages (Supplementary Fig. 2b-c). However, infiltrating macrophages in DA.VRA4 rats had lower expression (relative MFI levels) of MHCII+ compared to DA for naïve (0.76±0.15 vs 1.0±0.094, p=0.010, 95% CI [-0.42, -0.073]), α-Syn at 4 weeks (0.76±0.061 vs 1.0±0.046, p<0.001, 95% CI [-0.42, -0.13]) and control at 8 weeks (0.75±0.15 vs 0.95±0.17, p=0.026, 95% CI [-0.39, -0.024]) (Fig. 2f, g).

The expression (relative MFI levels) of CD86 (also known as B7-2, a co-stimulatory signal expressed by antigen-presenting cells necessary for activation of T lymphocytes [8]) on microglia and macrophages in the right/ipsilateral hemisphere did not differ between strains (Supplementary Fig. 2d-e). There were, however, lower CD86 levels in DA α-Syn vs control at 4 weeks (0.92±0.12 vs 1.2±0.08, p=0.004, 95% CI [-0.41, -0.08]) (Supplementary fig. 2e). No α-Syn- or strain-dependent differences were observed in percentages of infiltrating T lymphocytes in right/ipsilateral hemispheres (Supplementary fig. 2a, f).

### MHCII+ expression on blood myeloid cells and amount of CD3+ lymphocytes are regulated by α-Syn and *Ciita*

To investigate changes in peripheral immune cell populations, we performed flow cytometry of blood collected 4- and 8-weeks post nigral injection (Supplementary Fig. 3a). No strain- or α-Syn-dependent changes in overall percentage of myeloid (CD45+CD11b+) cells (Fig. 3a and Supplementary Fig. 3b) or MHCII+ myeloid cells were observed, except from more MHCII+ myeloid cells in naïve DA.VRA4 compared to naïve DA (12±2.6% vs 7.0±2.9%, p=0.0071, 95% CI [1.8, 8.9]) (Fig. 3b). Similar to MHCII+ macrophages in the brain, blood macrophages in DA.VRA4 rats had lower intensity of MHCII signal compared to DA α-Syn group at 4 weeks (1970±620 vs 2780±470 MFI, p=0.019, 95% CI [-1510, -100]) (Fig. 3c). Unlike the early response in microglia, blood myeloid cells from both strains displayed a peak MHCII+ response to α-Syn at 8 weeks in both strains (DA: 3230±1250 vs 2070±660 MFI, p=0.035, 95% CI [80, 2250] and DA.VRA4: 2640±350 vs 1480±250 MFI, p=0.021, 95% CI [170, 2150]) (Fig. 3c). We found no *Ciita*- or α-Syn-dependent regulation of CD86 expression on blood myeloid cells (Supplementary Fig. 3c).

**Fig. 3.**
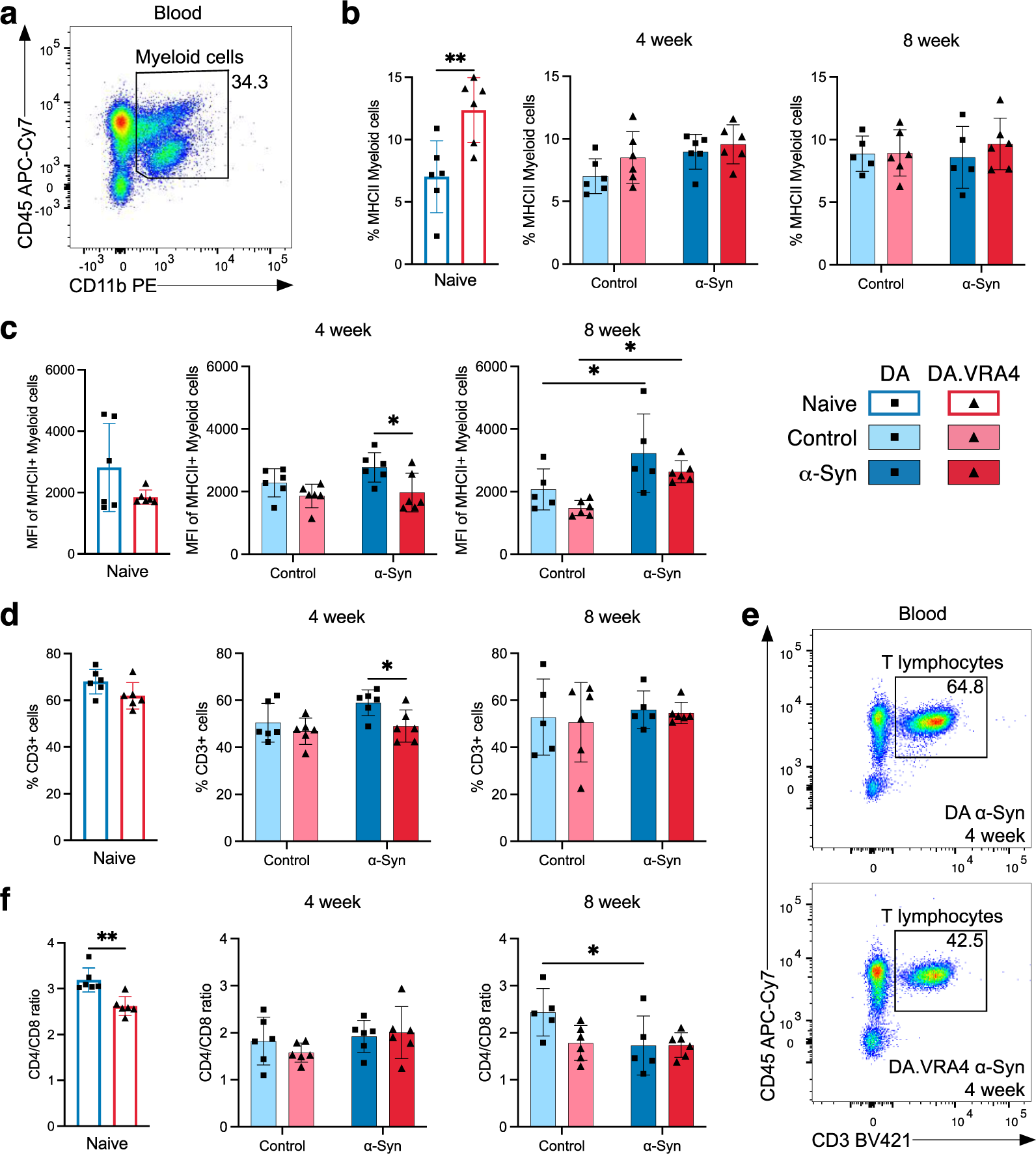
Systemic effects of rAAV-α-Syn+PFF on blood myeloid cells’ MHCII expression and circulating lymphocyte profiles. **a.** Gating of myeloid cells (CD45+CD11b+) in blood. **b.** Quantification of MHCII+ myeloid cells in blood. **c.** Quantification of MFI-values of MHCII+ myeloid cells in blood. **d.** Quantification of T lymphocytes (CD3+CD45+) in blood. **e.** Representative flow cytometric dot plot of circulating T lymphocyte population in DA (top) and DA.VRA4 (bottom) at 4 weeks in the α-Syn group. **f.** Quantification of CD4/CD8 ratio in blood. **b-d and f.** Naïve (DA n=6, DA.VRA4 n=6), 4 week; control (DA n=6, DA.VRA4 n=6) and α-Syn (DA n=6, DA.VRA4 n=6), 8 week; control (DA n=5, DA.VRA4 n=6) and α-Syn (DA n=5, DA.VRA4 n=6). α-Syn=rAAV-α-Syn+PFF. Control=rAAV-(-)+DPBS. Naïve rats were compared by unpaired Student’s t-test. Groups at 4- and 8-weeks were compared with two-way ANOVA with Šídák multiple comparison test (DA vs. DA.VRA4 and control vs. α-Syn). * p < 0.05 and ** p < 0.01. Data presented as mean ± SD with individual values.

The overall percentage of circulating T lymphocytes was lower in DA.VRA4 compared to DA rats in the α-Syn group at 4 weeks (49±6.8% vs 59±5.6%, p=0.021, 95% CI [-18, -1.9]) but did not differ at 8 weeks or between α-Syn and control groups (Fig. 3d, e). Among CD3+ lymphocytes, the CD4/CD8 ratio was lower in naïve DA.VRA4 compared to DA (2.6±0.21 vs 3.2±0.26, p=0.0019, 95% CI [-0.87, -0.27]) (Fig. 3f) which was driven by a decrease in CD4+ T lymphocytes (70±2.5% vs 74±1.4%, p=0.0037, 95% CI [-7.0, -1.8]) (Supplementary Fig. 3d) and an increase in CD8+ T lymphocytes (27±1.5% vs 23±1.4%, p=0.0023, 95% CI [1.5, 5.2]) (Supplementary Fig. 3e). Additionally, the CD4/CD8 ratio was reduced in response to α-Syn compared to control in DA rats at 8 weeks (1.7±0.63 vs 2.4±0.50, p=0.045, 95% CI [-1.4, -0.01]) (Fig. 3f), driven by an increase of CD8+ T lymphocytes (37±7.3% vs 28±4.9%, p=0.047, 95% CI [0.16, 18]) (Supplementary Fig. 3e).

### Few changes in CSF and serum cytokines but consistently higher levels of serum TNF in DA.VRA4 rats with lower levels of *Ciita* compared to DA

Altered levels of cytokines in the CSF and serum have been reported in PD patients [3]. To investigate the effect of *Ciita* expression and rAAV-α-Syn+PFF injection on CSF and serum cytokine levels, we performed multiplexed ELISA. Most of the cytokines investigated were detectable in both CSF and serum samples, however, the majority of the cytokines were below the lower limit of quantification, ultimately affecting the accuracy of the results (results summarized in Supplementary table 1-6).

Most cytokines analyzed in CSF levels were unaffected by *Ciita* and α-Syn, but we found differences in IL-10 and IL-6. In the α-Syn group at 8 weeks, DA.VRA4 rats had higher IL-10 CSF levels compared to DA (3.2±1.1 pg/ml vs 1.7±0.74 pg/ml, p=0.0084, 95% CI [0.42, 2.4]) (Fig. 4a). rAAV-α-Syn+PFF induced increased CSF levels of IL-6 at 8 weeks in both DA (45±21 pg/ml vs 12 ± 12 pg/ml, p<0.001, 95% CI [16, 50]) and DA.VAR4 (31±11 pg/ml vs 6.6 ± 7.6 pg/ml, p=0.003, 95% CI [8.1, 41]) (Fig.4b)

**Fig. 4.**
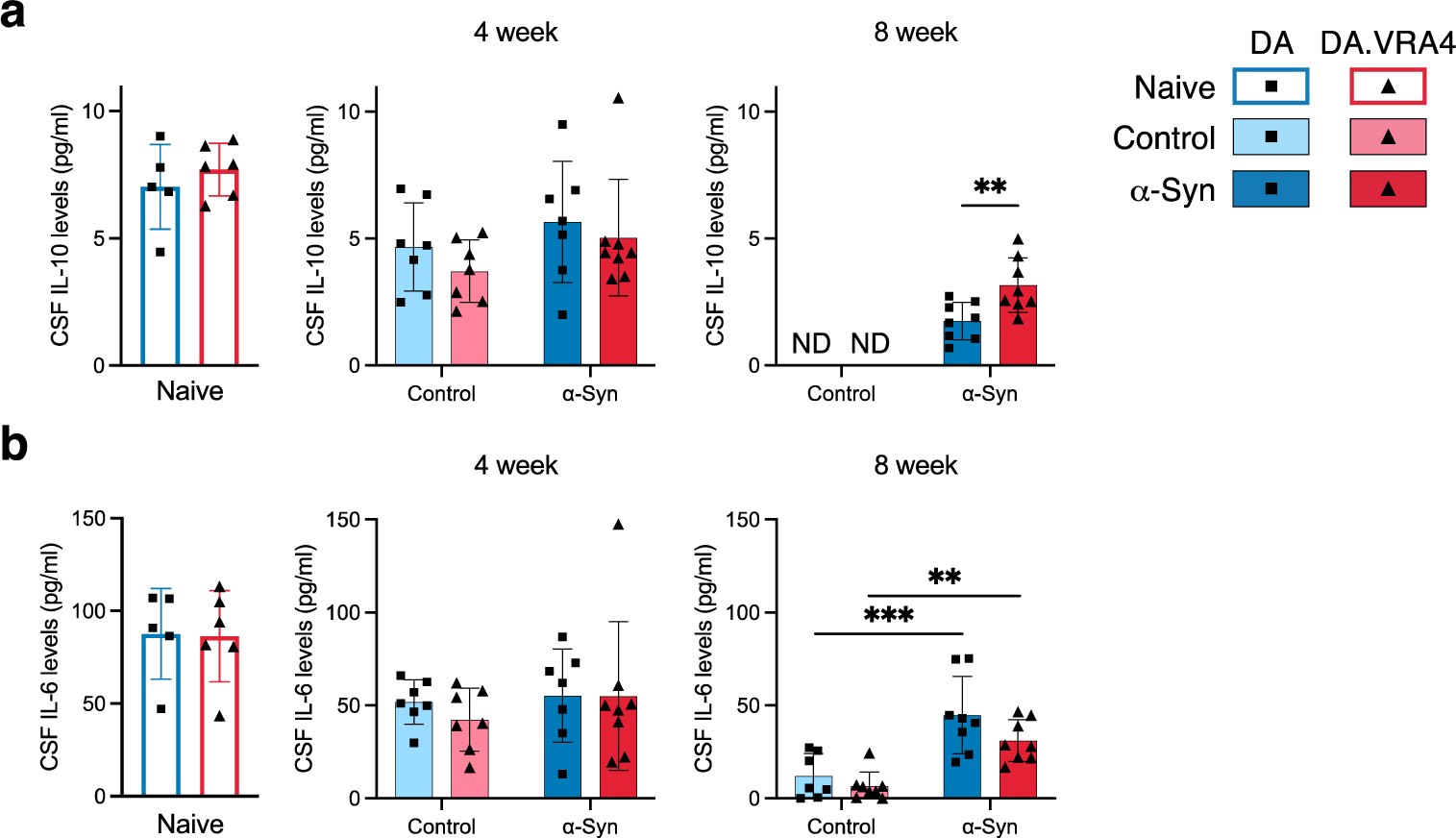
CSF IL-10 and IL-6 cytokine levels are regulated by differing *Ciita* levels and in response to rAAV-α-Syn+PFF, respectively. **a.** Quantification of IL-10 levels in cerebrospinal fluid (CSF) measured by ELISA. Non-detected levels indicated by “ND”. **b.** Quantification of IL-6 levels in CSF measured by ELISA. **a-b.** Naïve (DA n=5, DA.VRA4 n=6), 4-week; control (DA n=7, DA.VRA4 n=7) and α-Syn (DA n=7, DA.VRA4 n=8), 8-week; control (DA n=7, DA.VRA4 n=8) and α-Syn (DA n=8, DA.VRA4 n=8). α-Syn=rAAV-α-Syn+PFF. Control=rAAV-(-)+DPBS. Naïve rats were compared by unpaired Student’s t-test. Groups at 4- and 8-weeks were compared with two-way ANOVA with Šídák multiple comparison test (DA vs. DA.VRA4 and control vs. α-Syn). ** p < 0.01 and *** p < 0.001. Data presented as mean ± SD with individual values.

In serum, we found differences in IL-1β, IL-5 and TNF. IL-1β levels were higher in naïve DA.VRA4 compared to DA (29±14 pg/ml vs 11±8.3 pg/ml, p=0.025, 95% CI: [2.8, 33]) and at 4 weeks in the α-Syn group in DA.VRA4 rats (23±6.7 pg/ml vs 14±5.8 pg/ml, p=0.022, 95% CI [1.5, 16]) (Fig. 5a). IL-5 levels were unaffected by α-Syn but were higher in DA.VRA4 compared to DA rats in the α-Syn group at 4 weeks (37±5.8 pg/ml vs 25±11 pg/ml, p=0.007, 95% CI [3.2, 22] (Fig. 5b).

**Fig. 5.**
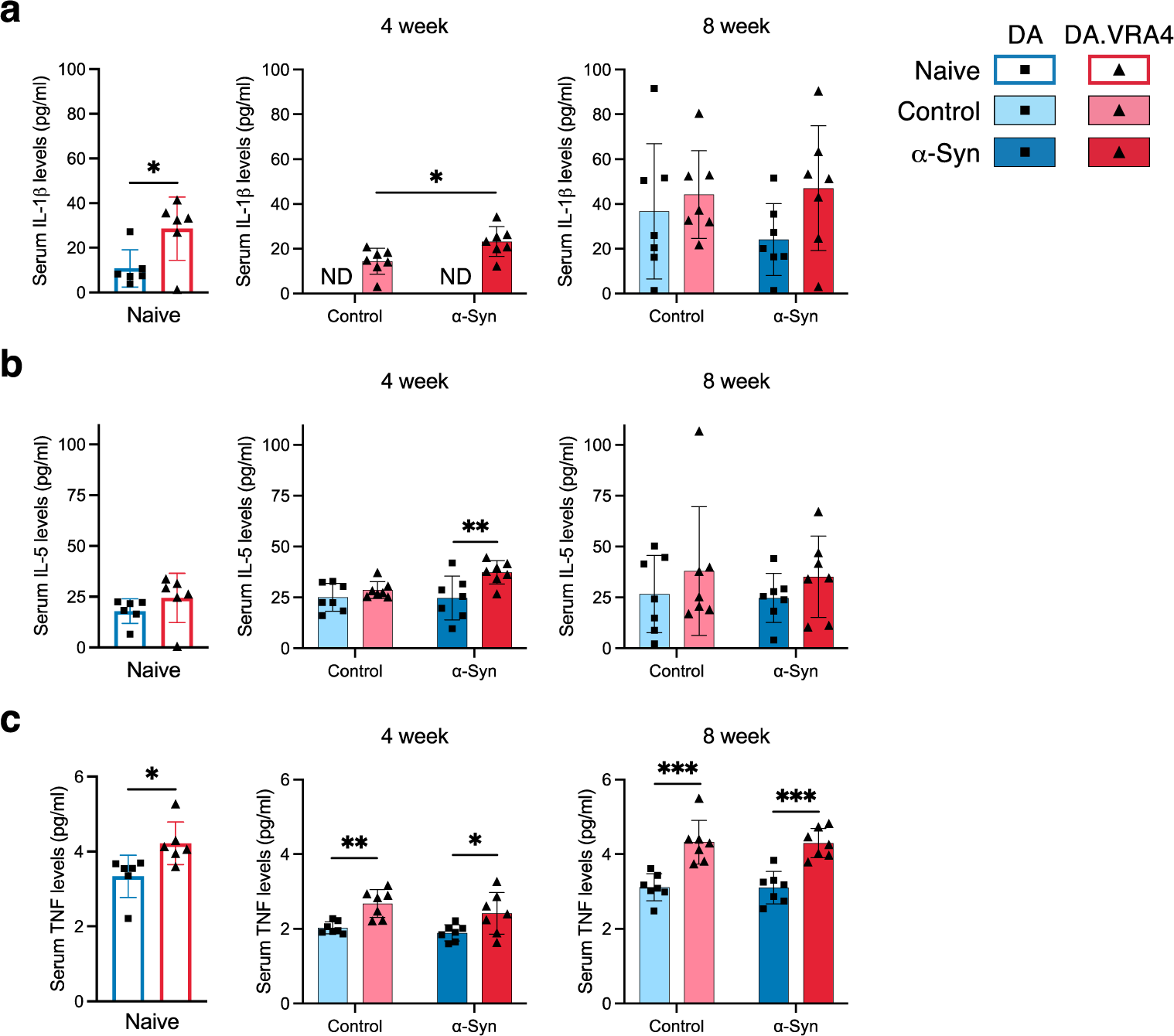
*Ciita* regulates TNF levels in serum. **a.** Quantification of serum IL-1β levels measured by ELISA. Non-detected levels indicated by “ND”. **b.** Quantification of serum IL-5 levels measured by ELISA. **c.** Quantification of TNF levels in serum measured by ELISA. **a-c.** Naïve (DA n=6, DA.VRA4 n=6), 4-week; control (DA n=7, DA.VRA4 n=7) and α-Syn (DA n=7, DA.VRA4 n=7), 8-week; control (DA n=7, DA.VRA4 n=7) and α-Syn (DA n=7, DA.VRA4 n=7). α-Syn=rAAV-α-Syn+PFF. Control=rAAV-(-)+DPBS. Naïve rats were compared by unpaired Student’s t-test. Groups at 4- and 8-weeks were compared with two-way ANOVA with Šídák multiple comparison test (DA vs. DA.VRA4 and control vs. α-Syn). * p <0.05, ** p < 0.01 and *** p < 0.001. Data presented as mean ± SD with individual values.

DA.VRA4 rats with lower levels of *Ciita* had consistently higher levels of TNF in serum compared to DA in naïve (4.2±0.57 pg/ml vs 3.3±0.56 pg/ml, p=0.022, 95% CI [0.15, 1.6]), control-(4-week: 2.7±0.37 pg/ml vs 2.0±0.16 pg/ml, p=0.0012, 95% CI [0.31, 0.98] and 8-week: 4.3±0.58 pg/ml vs 3.1±0.36 pg/ml, p=0.00060, 95% CI [0.64, 1.8]) and α-Syn groups (4-week: 2.4±0.56 pg/ml vs 1.9±0.22 pg/ml, p=0.039, 95% CI [0.030, 1.0] and 8-week: 4.3±0.40 pg/ml vs 3.1±0.43 pg/ml, p=0.00020, 95% CI [0.71, 1.7]) (Fig. 5c).

## DISCUSSION

Studies investigating human cohorts and experimental models support a role for antigen presentation and adaptive immune responses in PD etiology. However, there are contradictory findings on how resident and peripheral immune responses contribute to or protect against neuropathology in PD. Contributing factors to these discrepancies likely include difficulties in determining causality versus consequence in an ongoing pathological process, as well as the multiple different murine models used to study PD-related changes in the immune system. The use of transgenic models to model complex immune responses in human disease can be questioned. Therefore, we used a congenic rat model with naturally occurring *Ciita* alleles mediating differential expression of *Ciita* and *MHC*-genes in both rats and humans [16]. The CIITA protein regulates MHCII expression, and is a crucial link between antigen-presenting cells in the innate immune system and lymphocytes in the adaptive immune system. In a recent study, we showed that lower *Ciita* levels in rats are associated with increased susceptibility to α-Syn pathology and dopaminergic neurodegeneration [14, 15]. This strongly supports CIITA, MHCII and the process of antigen presentation to have causal impact on PD risk and outcome. The relative contribution of resident (brain) and peripheral (systemic) immune cells and cytokines in this process is, however, not known. Therefore, we characterized the effects of *Ciita* levels on local and peripheral immune populations and cytokines in the rAAV-α-Syn+PFF rat PD model. We used the DA.VRA4 strain with variants in the *Ciita* gene mediating lower expression of MHCII and the background strain, DA. These congenic rats provide a physiologically highly relevant model to study the effects of antigen presentation on immune populations and PD-like pathology, especially since genetic variants in the human orthologue *CIITA* also regulate MHCII expression and are associated with susceptibility to rheumatoid arthritis, multiple sclerosis and myocardial infarction [16].

Results from different PD models confirm an influence of MHCII on PD-like pathology, but are contradictory in terms of the direction of effect from altered *Ciita* and *MHCII* expression. This could be explained by the heterogeneity in study design, including the use of different species (rats [14, 15, 24] or mice [13, 25, 26]), different models of PD/synucleinopathies (transgenic [25], rAAV-α-Syn [13, 14, 24, 26], or rAAV-α-Syn+PFF [15]) and different immunological models (knock-out (KO) models [13, 25, 26], nude rats [24], silencing through shRNA [13], or congenic strains [14, 15]). In response to α-Syn overexpression, *Mhc2*-KO, *Ciita*-KO and *Ciita* silencing protected mice against dopaminergic cell loss [13, 26] and led to reduction of T lymphocyte- and monocyte infiltration [13], microglial activation [26] and amount of MHCII+ microglia (CD45^dim^CD11b+) [13]. In contrast, *Mhc2*-KO in mice expressing human α-Syn with the A53T mutation (M83+/0) resulted in accelerated pathology in the brain and an overall reduction of T lymphocytes in the CNS after injection of PFF into the hindlimb [25]. Our data presented here and previously [14, 15] also indicate lower *Ciita* levels to increase susceptibility to PD-like pathology. However, the nature of the respective *in vivo* models is important to consider when interpreting these results. In humans, KO of *CIITA* or other crucial transcription factors for MHCII causes severe immunodeficiency (bare lymphocyte syndrome, BLS). It is therefore likely that *Ciita*- and *Mhc2*-KO models have compromised immune systems that limit their physiological relevance. Silencing of *Ciita* circumvents this issue, and might be the best model to study effects of immunotherapies, while our congenic model comes closest to the situation in humans where common *CIITA* variants regulate MHCII expression and disease susceptibility throughout life, including myeloid- and lymphocyte development and thymic selection. Of note, previous results on genetic association between *CIITA* and risk for multiple sclerosis, rheumatoid arthritis and myocardial infarction show a similar direction of effect as found in our congenic rat model, with the risk allele in *CIITA* being associated with lower expression of *CIITA, CD74, HLA-DRA* and *HLA-DQA1* [16]. The combined genetic- and disease model employed in the current study has high construct validity (common *CIITA* genetic variants regulate MHCII levels both in rats and humans) and high face validity (the rAAV-α-Syn+PFF rat model displays seeded α-Syn pathology, dopaminergic neurodegeneration, motor impairment and neuroinflammation). Together, these characteristics allow for a good predictive validity of the model.

Using flow cytometry, we show that differential expression of *Ciita* affects the baseline levels of microglia and microglial MHCII-expression in naïve rats. Additionally, this study confirms previous semi-quantitative findings from brain immunostaining and RT-qPCR regarding microglial MHCII expression in response to α-Syn; lower *Ciita* levels in DA.VRA4 rats are associated lower levels of MHCII per cell [14, 15]. We hypothesize that increased numbers of MHCII+ microglia in naïve rats and region-specific differences in response to α-Syn (more MHCII+ microglia in striatum [15]) affect susceptibility and accelerate dopaminergic neurodegeneration through pathological spread of α-Syn (Fig. 6), as we have previously reported an increased aggregation and propagation of α-Syn in DA.VRA4 rats, along with pathological α-Syn (pS129) co-localized within MHCII+ microglial cells in the rAAV-α-Syn+PFF model [15]. In contrast to our hypothesized model a recent study suggests that the main contributor to α-Syn-induced neurodegeneration is border-associated macrophages (BAMs) rather than microglia [27]. In the current study we were unable to investigate any specific contribution of infiltrating BAMs in the brain parenchyma since they are indistinguishable from microglia based on CD45- and CD11b-expression [28].

**Fig. 6.**
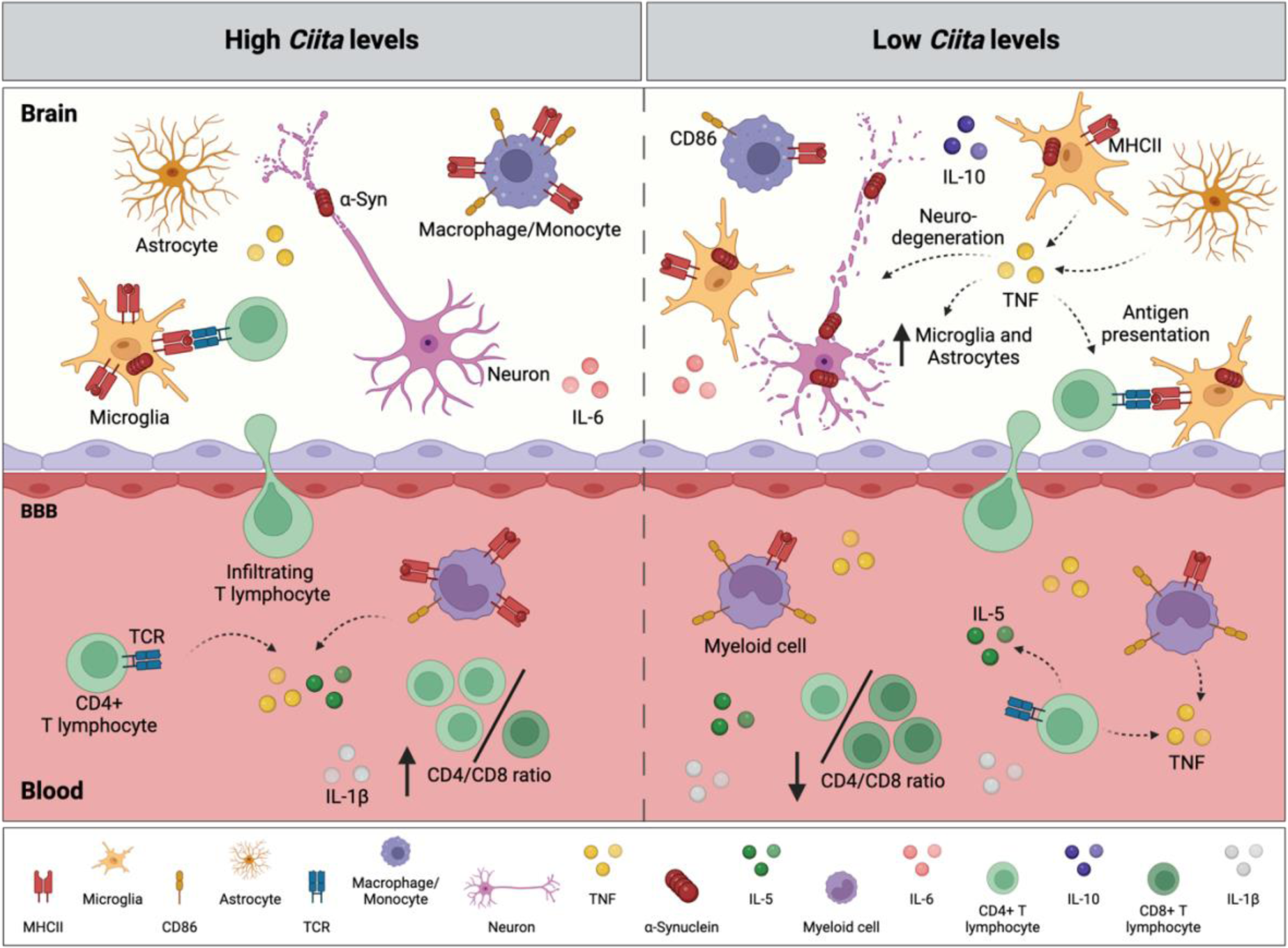
Hypothesis for how *Ciita* regulates baseline immune populations, cytokine levels, α-Syn propagation and susceptibility to PD-like neurodegeneration. Lower *Ciita* levels are associated with motor impairments [14, 15], neurodegeneration [14, 15] and exacerbated α-Syn pathological spread [15] in response to α-Syn. In this study, we found that lower *Ciita* levels are associated with lower MHCII levels on microglia [14, 15] and on myeloid cells in both brain and blood. Histology has shown an increased number of activated microglia in striatum [14, 15], and the current study shows that reduced *Ciita* levels are associated with increased numbers of MHCII+ microglia in brain and MHCII+ myeloid cells in circulation in naïve rats, with lower amount of MHCII per cell. We also found as higher cytokine (IL-5, TNF) levels in serum and a lower CD4/CD8 ratio in blood. In the proposed scenario, processed α-Syn peptides are presented to brain-infiltrating T lymphocytes, initiating an adaptive immune response. TNF could increase the number of nigral astrocytes and microglia as well as dopaminergic neurodegeneration, as has been shown in toxin- or endotoxin-induced PD models [49, 51, 52]. Dashed arrows indicate possible sources and/or mechanisms based on literature. Illustration created with BioRender.com.

Compared to studies using α-Syn nigral overexpression or striatal PFF injection in mice [13, 29, 30], we found very limited numbers of brain-infiltrating macrophages/monocytes and lymphocytes in the rAAV-α-Syn+PFF rat model. Among live cells analyzed from brain tissue, 93-97% were CD45^dim^CD11b+ microglia and only 0.5-1.5% CD45^high^CD11b+ macrophages/monocytes. However, a large majority (70-85%) of macrophages/monocytes but a minority (5-15%) of the microglia were MHCII+, indicating that infiltrating macrophages/monocytes might still play an important role in CNS antigen presentation. The low number of infiltrating macrophages/monocytes and lack of differences between α-Syn and control groups in this model are contradictory to a previous study in a nigral α-Syn overexpression model in mice, where PD-like pathology was mainly driven by infiltrating monocytes [31]. In addition, *Ciita* levels did not affect the number of infiltrating macrophages/monocytes or lymphocytes in our model, while KO and silencing of *Ciita* have been reported to greatly reduce both monocyte and lymphocyte infiltration in mice overexpressing α-Syn in SN [13]. Additionally, we show that naïve DA.VRA4 rats with lower *Ciita* levels than DA also have a higher percentage of MHCII+ circulating cells of the myeloid lineage and that these circulating myeloid cells have lower MHCII expression compared to DA in the α-Syn group at 4 weeks. The MHCII expression on circulating myeloid cells also peaked at α-Syn 8 weeks in both DA.VRA4 and DA.

We found few infiltrating T lymphocytes (CD45+CD3+) in brain tissue, and the amounts were independent of *Ciita* levels. In blood, DA.VRA4 rats with lower *Ciita* levels had fewer T lymphocytes compared to DA at 4 weeks after rAAV-α-Syn+PFF. There was also a lower CD4/CD8 ratio in naïve DA.VRA4 rats compared to DA. Studies in other α-Syn-based PD models indicate both detrimental and protective roles of lymphocytes. Neurodegeneration-promoting effects are supported by findings that mice lacking lymphocytes (*Rag1* KO) were protected against dopaminergic cell loss in SN, that lymphocyte reconstitution resulted in dopaminergic cell loss comparable to wt mice [32] and that CD4 KO protected against neurodegeneration in the SN and inhibited myeloid activation [33]. In contrast, protective effects of lymphocytes have been reported in a striatal PFF model, where adoptive transfer of CD4+ lymphocytes to immunocompromised mice reduced α-Syn pathology [34]. More detailed studies are required to delineate if the observed differences in the current study are of biological relevance to susceptibility and/or progression of PD-like pathology.

Although recent genetic studies point at MHCII, but not MHCI, genes associated to PD [7], neuronal MHCI expression [35], CD8+ T lymphocyte infiltration [36] and reactivity to α-Syn [11] could potentially play a role in PD susceptibility. Studies on lymphocyte populations in PD patients are, however, inconclusive. Different studies have reported no difference [37] or decrease [38] in CD4+ and CD8+ populations, a reduction in effector and regulatory T lymphocytes [37], a lower CD4/CD8 ratio [39] and an overall decrease in circulating CD4+ lymphocyte subpopulations due to decreased levels of T-helper (Th) 2, Th17 and regulatory lymphocytes [40]. Other research suggests an overall decrease in circulating lymphocytes with increased Th1 and Th17 but decreased Th2 and regulatory T lymphocytes [41] or no changes in Th1 and Th2 subsets but an increase in the Th17 lymphocyte population [42]. In addition to differences in population sizes and ratios, functional studies indicate alterations in lymphocyte populations in PD, including deficits in migratory capacity of CD4+ T lymphocytes from PD patients [43] and impaired suppressor functions of T regulatory cells in PD, which could be restored by *ex vivo* expansion [37]. A third study reported that PD disease severity was associated with higher activation levels of T lymphocytes in response to phytohemagglutinin stimulation [38]. In light of these findings from PD patients, a limitation of our study is not assessing *Ciita* effects on MHCI expression, another is the low number of lymphocytes detected in brain tissue that limited analyses of sub-populations of infiltrating T lymphocytes.

In addition to altered immune cell profiles, we analyzed CSF and serum levels of cytokines. Although few differences were found, there are several links to PD for the cytokines that differed depending on strain and/or rAAV-α-Syn+PFF injection. Elevated IL-6 levels in CSF have been observed in PD patients [44] and we found that rAAV-α-syn+PFF injection resulted in increased CSF IL-6 levels in both DA and DA.VRA4 rats. The anti-inflammatory cytokine IL-10 has previously been shown to be neuroprotective and reduce microglial activation in toxin models of PD [45]. We found higher IL-10 levels in CSF from DA.VRA4 rats compared to DA in response to α-Syn at 8-weeks despite the more severe neuropathological phenotype observed in DA.VRA4 rats [15]. IL-1β levels in blood have been reported in multiple studies to be increased in PD patients [46] and to correlate with disease progression [47]. IL-1β levels have also been shown to influence the NLRP3 inflammasome which contributes to neurodegeneration in a 6-OHDA mouse model of PD [48]. We found higher levels of IL-1β in naïve DA.VRA4 vs. DA and a significant upregulation in DA.VRA4 rats in response to α-Syn. IL-5 has been reported to be elevated in CD4+ Th2 lymphocytes from PD patients stimulated with α-Syn peptides *ex vivo* [11]. We found higher IL-5 levels in serum from DA.VRA4 compared to DA in the α-Syn group at 4 weeks.

Interestingly, we found consistently higher TNF levels in serum but not CSF in DA.VRA4 compared to DA rats. Other studies have shown that intrathecal neutralization of soluble TNF by dominant-negative protein (DN-TNF) in SN, but not striatum, protects from 6-OHDA- and LPS-induced degeneration of rat nigral dopaminergic neurons and neuroinflammation *in vivo* and *in vitro* [49, 50]. The neuroprotective effect of viral vector-mediated expression of DN-TNF was still present when administered 2 weeks after 6-OHDA lesion [51], while peripheral administration of DN-TNF was shown to cross the blood-brain barrier, reduce astrocyte- and microglial number in SN and to reduce neurodegeneration when administered 3 days, but not 14 days, after striatal 6-OHDA lesion [52]. Based on this data, TNF could exert direct and indirect effects on neurons, astrocytes, microglia and T lymphocytes. Further, it is possible that higher levels of TNF in DA.VRA4 rats affect the susceptibility to PD-like pathology and together with IL-1β and IL-5 exacerbates α-Syn pathological spread and dopaminergic neurodegeneration (Fig. 6). Further investigation is necessary to provide a mechanistic view, i.e. by assessing the effects of systemic TNF inhibition on neuroinflammation, neurodegeneration and α-Syn pathology in the rAAV-α-syn+PFF model in the DA.VRA4 rats with elevated TNF serum levels.

As all models, the rAAV-α-Syn+PFF PD rat model has both strengths and limitations. Similar to other models using intracranial injections, the physical damage and blood-brain barrier disruption could cause changes in immune populations, independent of what is injected. In order to control for immune responses not related to α-Syn, we used an empty vector in SN and vehicle in striatum for the control groups. We chose an empty vector since we and others have observed that the commonly used rAAV-GFP control vector elicits a neuroinflammatory response [15, 31]. As control for PFF, we chose to use vehicle, since we have found that bovine serum albumin elicits a neuroinflammatory response [15] and other studies report that α-Syn monomers and saline are comparable controls for the PFF model in rats [53]. This design, however, means that we cannot determine to what extent the expression of a foreign protein or possible contamination with endotoxin in the PFF preparation affect the results. Another aspect to consider is the timepoints selected. We have previously investigated the effects of differential *Ciita* expression on PD-like α-Syn pathology, neurodegeneration and neuroinflammation at 8-weeks in the rAAV-α-Syn+PFF model [15] and at 12-weeks in the rAAV-α-Syn model [14]. However, studies have shown that there is an inflammatory response ongoing prior to neurodegeneration in animal models [30, 53, 54] and in PD patients [11]. We chose to include two timepoints (4- and 8 weeks post nigral injection), but acknowledge that the results might differ at other timepoints. A technical limitation is that we used entire hemispheres for flow cytometric analyses of brain tissue. Although α-Syn pathology and MHCII+ microglial cells are widespread in the brain in the rAAV-α-Syn+PFF model [15], it is possible that region-specific differences affected by *Ciita* levels or responses to α-Syn are larger than what was reflected in entire hemispheres.

Even though there is substantial evidence of the involvement of antigen presentation in PD based on genetic association studies and elevated MHCII levels at the site of neurodegeneration [3-5, 7, 55, 56], sufficient knowledge on the role of MHCII in disease etiology is lacking. Association between HLA alleles and PD risk has been found for expression quantitative trait loci (eQTL) [4-6], non-coding variants [7, 57] and non-synonymous coding variants [7]. The heterogeneity and strong linkage within the HLA make discrimination between these effects difficult. In light of previous findings and the current study that point towards *Ciita*-mediated effects on PD-like pathology, we hypothesize that alleles affecting *CIITA* expression interact with non-coding risk-HLA alleles affecting *MHCII* expression (eQTLs). Further, *CIITA* expression could modify the effect of coding risk-HLA alleles by affecting their expression levels. In addition, HLA alleles have been reported to interact with other non-genetic factors, including pyrethroids and smoking [5, 58] and *CIITA* could potentially be part of such interactive effects.

In conclusion, despite significant differences in PD-relevant neurodegenerative and behavioral phenotypes, we observed only subtle differences in immune cell populations and cytokine profiles between the more susceptible DA.VRA4 and the more resistant DA rats, the most consistent being higher levels of serum TNF in DA.VRA4. To determine if and how these immune effects are causally related to the increased susceptibility to α-Syn-induced PD-like pathology, further studies are required, e.g., with specific cytokine inhibitors. Our work together with other experimental and human studies highlight the complexity and importance of understanding the link between innate and adaptive immune responses in PD. It also suggests that future immunomodulatory therapies for PD could be efficient without major impact on the immune system as a whole.

## ACKNOWLEDGEMENTS

We want to acknowledge the AAV Vector Lab platform and Jenny G. Johansson for production of the rAAV constructs, the FACS Platform and Anna Hammarberg for assistance of flow cytometry experiments, the MESO QuickPlex Platform and Shorena Janelidze for access to the MESO QuickPlex SQ 120 analyzer, the Confocal Microscope Platform for access to the Leica SP8 scanning confocal microscope, Lund University Bioimaging Centre (LBIC) for providing experimental resources and Lund Stem Cell Center Imaging Facility for access to the Olympus VS-120 virtual slide microscope.

## DATA AVAILABILITY

The data supporting the findings of this study are available on request from the corresponding author.

## REFERENCES

[1] Poewe W, Seppi K, Tanner CM, Halliday GM, Brundin P, Volkmann J, Schrag AE, Lang AE (2017) Parkinson disease. Nat Rev Dis Primers 3, 17013.

[2] Ascherio A, Schwarzschild MA (2016) The epidemiology of Parkinson’s disease: risk factors and prevention. Lancet Neurol 15, 1257–1272.

[3] Hirsch EC, Hunot S (2009) Neuroinflammation in Parkinson’s disease: a target for neuroprotection? Lancet Neurol 8, 382–397.

[4] Hamza TH, Zabetian CP, Tenesa A, Laederach A, Montimurro J, Yearout D, Kay DM, Doheny KF, Paschall J, Pugh E, Kusel VI, Collura R, Roberts J, Griffith A, Samii A, Scott WK, Nutt J, Factor SA, Payami H (2010) Common genetic variation in the HLA region is associated with late-onset sporadic Parkinson’s disease. Nat Genet 42, 781–785.

[5] Kannarkat GT, Cook DA, Lee JK, Chang J, Chung J, Sandy E, Paul KC, Ritz B, Bronstein J, Factor SA, Boss JM, Tansey MG (2015) Common Genetic Variant Association with Altered HLA Expression, Synergy with Pyrethroid Exposure, and Risk for Parkinson’s Disease: An Observational and Case-Control Study. NPJ Parkinsons Dis 1.

[6] Wissemann WT, Hill-Burns EM, Zabetian CP, Factor SA, Patsopoulos N, Hoglund B, Holcomb C, Donahue RJ, Thomson G, Erlich H, Payami H (2013) Association of Parkinson disease with structural and regulatory variants in the HLA region. Am J Hum Genet 93, 984–993.

[7] Yu E, Ambati A, Andersen MS, Krohn L, Estiar MA, Saini P, Senkevich K, Sosero YL, Sreelatha AAK, Ruskey JA, Asayesh F, Spiegelman D, Toft M, Viken MK, Sharma M, Blauwendraat C, Pihlstrom L, Mignot E, Gan-Or Z (2021) Fine mapping of the HLA locus in Parkinson’s disease in Europeans. NPJ Parkinsons Dis 7, 84.

[8] Huppa JB, Davis MM (2003) T-cell-antigen recognition and the immunological synapse. Nat Rev Immunol 3, 973–983.

[9] Brochard V, Combadiere B, Prigent A, Laouar Y, Perrin A, Beray-Berthat V, Bonduelle O, Alvarez-Fischer D, Callebert J, Launay JM, Duyckaerts C, Flavell RA, Hirsch EC, Hunot S (2009) Infiltration of CD4+ lymphocytes into the brain contributes to neurodegeneration in a mouse model of Parkinson disease. J Clin Invest 119, 182–192.

[10] Gate D, Tapp E, Leventhal O, Shahid M, Nonninger TJ, Yang AC, Strempfl K, Unger MS, Fehlmann T, Oh H, Channappa D, Henderson VW, Keller A, Aigner L, Galasko DR, Davis MM, Poston KL, Wyss-Coray T (2021) CD4(+) T cells contribute to neurodegeneration in Lewy body dementia. Science 374, 868–874.

[11] Sulzer D, Alcalay RN, Garretti F, Cote L, Kanter E, Agin-Liebes J, Liong C, McMurtrey C, Hildebrand WH, Mao X, Dawson VL, Dawson TM, Oseroff C, Pham J, Sidney J, Dillon MB, Carpenter C, Weiskopf D, Phillips E, Mallal S, Peters B, Frazier A, Lindestam Arlehamn CS, Sette A (2017) T cells from patients with Parkinson’s disease recognize alpha-synuclein peptides. Nature 546, 656–661.

[12] Lindestam Arlehamn CS, Dhanwani R, Pham J, Kuan R, Frazier A, Rezende Dutra J, Phillips E, Mallal S, Roederer M, Marder KS, Amara AW, Standaert DG, Goldman JG, Litvan I, Peters B, Sulzer D, Sette A (2020) alpha-Synuclein-specific T cell reactivity is associated with preclinical and early Parkinson’s disease. Nat Commun 11, 1875.

[13] Williams GP, Schonhoff AM, Jurkuvenaite A, Thome AD, Standaert DG, Harms AS (2018) Targeting of the class II transactivator attenuates inflammation and neurodegeneration in an alpha-synuclein model of Parkinson’s disease. J Neuroinflammation 15, 244.

[14] Jimenez-Ferrer I, Jewett M, Tontanahal A, Romero-Ramos M, Swanberg M (2017) Allelic difference in Mhc2ta confers altered microglial activation and susceptibility to alpha-synuclein-induced dopaminergic neurodegeneration. Neurobiol Dis 106, 279–290.

[15] Jimenez-Ferrer I, Backstrom F, Duenas-Rey A, Jewett M, Boza-Serrano A, Luk KC, Deierborg T, Swanberg M (2021) The MHC class II transactivator modulates seeded alpha-synuclein pathology and dopaminergic neurodegeneration in an in vivo rat model of Parkinson’s disease. Brain Behav Immun 91, 369–382.

[16] Swanberg M, Lidman O, Padyukov L, Eriksson P, Akesson E, Jagodic M, Lobell A, Khademi M, Borjesson O, Lindgren CM, Lundman P, Brookes AJ, Kere J, Luthman H, Alfredsson L, Hillert J, Klareskog L, Hamsten A, Piehl F, Olsson T (2005) MHC2TA is associated with differential MHC molecule expression and susceptibility to rheumatoid arthritis, multiple sclerosis and myocardial infarction. Nat Genet 37, 486–494.

[17] Thakur P, Breger LS, Lundblad M, Wan OW, Mattsson B, Luk KC, Lee VMY, Trojanowski JQ, Bjorklund A (2017) Modeling Parkinson’s disease pathology by combination of fibril seeds and alpha-synuclein overexpression in the rat brain. Proc Natl Acad Sci U S A 114, E8284–E8293.

[18] Decressac M, Mattsson B, Lundblad M, Weikop P, Bjorklund A (2012) Progressive neurodegenerative and behavioural changes induced by AAV-mediated overexpression of alpha-synuclein in midbrain dopamine neurons. Neurobiol Dis 45, 939–953.

[19] Harnesk K, Swanberg M, Ockinger J, Diez M, Lidman O, Wallstrom E, Lobell A, Olsson T, Piehl F (2008) Vra4 congenic rats with allelic differences in the class II transactivator gene display altered susceptibility to experimental autoimmune encephalomyelitis. J Immunol 180, 3289–3296.

[20] Luk KC, Song C, O’Brien P, Stieber A, Branch JR, Brunden KR, Trojanowski JQ, Lee VM (2009) Exogenous alpha-synuclein fibrils seed the formation of Lewy body-like intracellular inclusions in cultured cells. Proc Natl Acad Sci U S A 106, 20051–20056.

[21] Paxinos G, Watson C (2014) Paxinos and Watson’s The Rat Brain in Stereotaxic Coordinates, Elsevier Academic Press, San Diego.

[22] Grabert K, Michoel T, Karavolos MH, Clohisey S, Baillie JK, Stevens MP, Freeman TC, Summers KM, McColl BW (2016) Microglial brain region-dependent diversity and selective regional sensitivities to aging. Nat Neurosci 19, 504–516.

[23] Grabert K, McColl BW (2018) Isolation and Phenotyping of Adult Mouse Microglial Cells. Methods Mol Biol 1784, 77–86.

[24] Subbarayan MS, Hudson C, Moss LD, Nash KR, Bickford PC (2020) T cell infiltration and upregulation of MHCII in microglia leads to accelerated neuronal loss in an alpha-synuclein rat model of Parkinson’s disease. J Neuroinflammation 17, 242.

[25] Gonzalez De La Cruz E, Vo Q, Moon K, McFarland KN, Weinrich M, Williams T, Giasson BI, Chakrabarty P (2022) MhcII Regulates Transmission of alpha-Synuclein-Seeded Pathology in Mice. Int J Mol Sci 23.

[26] Harms AS, Cao S, Rowse AL, Thome AD, Li X, Mangieri LR, Cron RQ, Shacka JJ, Raman C, Standaert DG (2013) MHCII is required for alpha-synuclein-induced activation of microglia, CD4 T cell proliferation, and dopaminergic neurodegeneration. J Neurosci 33, 9592-9600.

[27] Schonhoff AM, Figge DA, Williams GP, Jurkuvenaite A, Gallups NJ, Childers GM, Webster JM, Standaert DG, Goldman JE, Harms AS (2023) Border-associated macrophages mediate the neuroinflammatory response in an alpha-synuclein model of Parkinson disease. Nat Commun 14, 3754.

[28] Mrdjen D, Pavlovic A, Hartmann FJ, Schreiner B, Utz SG, Leung BP, Lelios I, Heppner FL, Kipnis J, Merkler D, Greter M, Becher B (2018) High-Dimensional Single-Cell Mapping of Central Nervous System Immune Cells Reveals Distinct Myeloid Subsets in Health, Aging, and Disease. Immunity 48, 380–395 e386.

[29] Basurco L, Abellanas MA, Ayerra L, Conde E, Vinueza-Gavilanes R, Luquin E, Vales A, Vilas A, Martin-Uriz PS, Tamayo I, Alonso MM, Hernaez M, Gonzalez-Aseguinolaza G, Clavero P, Mengual E, Arrasate M, Hervas-Stubbs S, Aymerich MS (2022) Microglia and astrocyte activation is region-dependent in the alpha-synuclein mouse model of Parkinson’s disease. Glia.

[30] Earls RH, Menees KB, Chung J, Barber J, Gutekunst CA, Hazim MG, Lee JK (2019) Intrastriatal injection of preformed alpha-synuclein fibrils alters central and peripheral immune cell profiles in non-transgenic mice. J Neuroinflammation 16, 250.

[31] Harms AS, Thome AD, Yan Z, Schonhoff AM, Williams GP, Li X, Liu Y, Qin H, Benveniste EN, Standaert DG (2018) Peripheral monocyte entry is required for alpha-Synuclein induced inflammation and Neurodegeneration in a model of Parkinson disease. Exp Neurol 300, 179–187.

[32] Karikari AA, McFleder RL, Ribechini E, Blum R, Bruttel V, Knorr S, Gehmeyr M, Volkmann J, Brotchie JM, Ahsan F, Haack B, Monoranu CM, Keber U, Yeghiazaryan R, Pagenstecher A, Heckel T, Bischler T, Wischhusen J, Koprich JB, Lutz MB, Ip CW (2022) Neurodegeneration by alpha-synuclein-specific T cells in AAV-A53T-alpha-synuclein Parkinson’s disease mice. Brain Behav Immun 101, 194–210.

[33] Williams GP, Schonhoff AM, Jurkuvenaite A, Gallups NJ, Standaert DG, Harms AS (2021) CD4 T cells mediate brain inflammation and neurodegeneration in a mouse model of Parkinson disease. Brain.

[34] George S, Tyson T, Rey NL, Sheridan R, Peelaerts W, Becker K, Schulz E, Meyerdirk L, Burmeister AR, von Linstow CU, Steiner JA, Galvis MLE, Ma J, Pospisilik JA, Labrie V, Brundin L, Brundin P (2021) T Cells Limit Accumulation of Aggregate Pathology Following Intrastriatal Injection of alpha-Synuclein Fibrils. J Parkinsons Dis 11, 585–603.

[35] Cebrian C, Zucca FA, Mauri P, Steinbeck JA, Studer L, Scherzer CR, Kanter E, Budhu S, Mandelbaum J, Vonsattel JP, Zecca L, Loike JD, Sulzer D (2014) MHC-I expression renders catecholaminergic neurons susceptible to T-cell-mediated degeneration. Nat Commun 5, 3633.

[36] Galiano-Landeira J, Torra A, Vila M, Bove J (2020) CD8 T cell nigral infiltration precedes synucleinopathy in early stages of Parkinson’s disease. Brain 143, 3717–3733.

[37] Thome AD, Atassi F, Wang J, Faridar A, Zhao W, Thonhoff JR, Beers DR, Lai EC, Appel SH (2021) Ex vivo expansion of dysfunctional regulatory T lymphocytes restores suppressive function in Parkinson’s disease. NPJ Parkinsons Dis 7, 41.

[38] Bhatia D, Grozdanov V, Ruf WP, Kassubek J, Ludolph AC, Weishaupt JH, Danzer KM (2021) T-cell dysregulation is associated with disease severity in Parkinson’s Disease. J Neuroinflammation 18, 250.

[39] Bas J, Calopa M, Mestre M, Mollevi DG, Cutillas B, Ambrosio S, Buendia E (2001) Lymphocyte populations in Parkinson’s disease and in rat models of parkinsonism. J Neuroimmunol 113, 146–152.

[40] Kustrimovic N, Comi C, Magistrelli L, Rasini E, Legnaro M, Bombelli R, Aleksic I, Blandini F, Minafra B, Riboldazzi G, Sturchio A, Mauri M, Bono G, Marino F, Cosentino M (2018) Parkinson’s disease patients have a complex phenotypic and functional Th1 bias: cross-sectional studies of CD4+ Th1/Th2/T17 and Treg in drug-naive and drug-treated patients. J Neuroinflammation 15, 205.

[41] Chen Y, Qi B, Xu W, Ma B, Li L, Chen Q, Qian W, Liu X, Qu H (2015) Clinical correlation of peripheral CD4+cell subsets, their imbalance and Parkinson’s disease. Mol Med Rep 12, 6105–6111.

[42] Sommer A, Marxreiter F, Krach F, Fadler T, Grosch J, Maroni M, Graef D, Eberhardt E, Riemenschneider MJ, Yeo GW, Kohl Z, Xiang W, Gage FH, Winkler J, Prots I, Winner B (2018) Th17 Lymphocytes Induce Neuronal Cell Death in a Human iPSC-Based Model of Parkinson’s Disease. Cell Stem Cell 23, 123–131 e126.

[43] Mamula D, Khosousi S, He Y, Lazarevic V, Svenningsson P (2022) Impaired migratory phenotype of CD4(+) T cells in Parkinson’s disease. NPJ Parkinsons Dis 8, 171.

[44] Chen X, Hu Y, Cao Z, Liu Q, Cheng Y (2018) Cerebrospinal Fluid Inflammatory Cytokine Aberrations in Alzheimer’s Disease, Parkinson’s Disease and Amyotrophic Lateral Sclerosis: A Systematic Review and Meta-Analysis. Front Immunol 9, 2122.

[45] Kwilasz AJ, Grace PM, Serbedzija P, Maier SF, Watkins LR (2015) The therapeutic potential of interleukin-10 in neuroimmune diseases. Neuropharmacology 96, 55–69.

[46] Qin XY, Zhang SP, Cao C, Loh YP, Cheng Y (2016) Aberrations in Peripheral Inflammatory Cytokine Levels in Parkinson Disease: A Systematic Review and Meta-analysis. JAMA Neurol 73, 1316–1324.

[47] Fan Z, Pan YT, Zhang ZY, Yang H, Yu SY, Zheng Y, Ma JH, Wang XM (2020) Systemic activation of NLRP3 inflammasome and plasma alpha-synuclein levels are correlated with motor severity and progression in Parkinson’s disease. J Neuroinflammation 17, 11.

[48] Gordon R, Albornoz EA, Christie DC, Langley MR, Kumar V, Mantovani S, Robertson AAB, Butler MS, Rowe DB, O’Neill LA, Kanthasamy AG, Schroder K, Cooper MA, Woodruff TM (2018) Inflammasome inhibition prevents alpha-synuclein pathology and dopaminergic neurodegeneration in mice. Sci Transl Med 10.

[49] McCoy MK, Martinez TN, Ruhn KA, Szymkowski DE, Smith CG, Botterman BR, Tansey KE, Tansey MG (2006) Blocking soluble tumor necrosis factor signaling with dominant-negative tumor necrosis factor inhibitor attenuates loss of dopaminergic neurons in models of Parkinson’s disease. J Neurosci 26, 9365–9375.

[50] McCoy MK, Ruhn KA, Martinez TN, McAlpine FE, Blesch A, Tansey MG (2008) Intranigral lentiviral delivery of dominant-negative TNF attenuates neurodegeneration and behavioral deficits in hemiparkinsonian rats. Mol Ther 16, 1572–1579.

[51] Harms AS, Barnum CJ, Ruhn KA, Varghese S, Trevino I, Blesch A, Tansey MG (2011) Delayed dominant-negative TNF gene therapy halts progressive loss of nigral dopaminergic neurons in a rat model of Parkinson’s disease. Mol Ther 19, 46–52.

[52] Barnum CJ, Chen X, Chung J, Chang J, Williams M, Grigoryan N, Tesi RJ, Tansey MG (2014) Peripheral administration of the selective inhibitor of soluble tumor necrosis factor (TNF) XPro(R)1595 attenuates nigral cell loss and glial activation in 6-OHDA hemiparkinsonian rats. J Parkinsons Dis 4, 349–360.

[53] Harms AS, Delic V, Thome AD, Bryant N, Liu Z, Chandra S, Jurkuvenaite A, West AB (2017) alpha-Synuclein fibrils recruit peripheral immune cells in the rat brain prior to neurodegeneration. Acta Neuropathol Commun 5, 85.

[54] Duffy MF, Collier TJ, Patterson JR, Kemp CJ, Luk KC, Tansey MG, Paumier KL, Kanaan NM, Fischer DL, Polinski NK, Barth OL, Howe JW, Vaikath NN, Majbour NK, El-Agnaf OMA, Sortwell CE (2018) Lewy body-like alpha-synuclein inclusions trigger reactive microgliosis prior to nigral degeneration. J Neuroinflammation 15, 129.

[55] McGeer PL, Itagaki S, Boyes BE, McGeer EG (1988) Reactive microglia are positive for HLA-DR in the substantia nigra of Parkinson’s and Alzheimer’s disease brains. Neurology 38, 1285–1291.

[56] Tansey MG, Wallings RL, Houser MC, Herrick MK, Keating CE, Joers V (2022) Inflammation and immune dysfunction in Parkinson disease. Nat Rev Immunol 22, 657–673.

[57] International Parkinson Disease Genomics C, Nalls MA, Plagnol V, Hernandez DG, Sharma M, Sheerin UM, Saad M, Simon-Sanchez J, Schulte C, Lesage S, Sveinbjornsdottir S, Stefansson K, Martinez M, Hardy J, Heutink P, Brice A, Gasser T, Singleton AB, Wood NW (2011) Imputation of sequence variants for identification of genetic risks for Parkinson’s disease: a meta-analysis of genome-wide association studies. Lancet 377, 641-649.

[58] Hollenbach JA, Norman PJ, Creary LE, Damotte V, Montero-Martin G, Caillier S, Anderson KM, Misra MK, Nemat-Gorgani N, Osoegawa K, Santaniello A, Renschen A, Marin WM, Dandekar R, Parham P, Tanner CM, Hauser SL, Fernandez-Vina M, Oksenberg JR (2019) A specific amino acid motif of HLA-DRB1 mediates risk and interacts with smoking history in Parkinson’s disease. Proc Natl Acad Sci U S A 116, 7419–7424.

