## Supplemental information for "MHC class II transactivator effects on local and systemic immune responses in an α-synuclein seeded rat model for Parkinson’s disease"

### SUPPLEMENTARY DATA

**Table S1.** Cytokine levels in CSF from naïve DA and DA.VRA4 rats. Lower limit of quantification (LLOQ) is specified in brackets after each cytokine. Data presented as mean±SD. ND=non-detected. Unpaired Student's t-test.

| CSF cytokines (pg/ml) | DA Naïve (n=5) | DA.VRA4 Naïve (n=6) |
|---|---|---|
| **IFNγ (39.7)** | 4.59±0.887 | 4.96±1.17 |
| **IL-10 (163)** | 7.02±1.67 | 7.70±1.04 |
| **IL-13 (12.5)** | ND | ND |
| **IL-1β (102)** | 16.1±6.22 | 15.5±3.26 |
| **IL-4 (8.00)** | 1.10±0.606 | 1.23±0.577 |
| **IL-5 (82.0)** | 37.6±12.3 | 36.5±11.5 |
| **IL-6 (96.9)** | 87.6±24.4 | 86.3±24.6 |
| **KC/GRO (21.7)** | 72.7±14.4 | 87.8±8.69 |
| **TNF (9.10)** | 1.48±0.337 | 1.57±0.306 |

**Table S2.** No differences in cytokine levels in CSF from DA and DA.VRA4 rats in the α-Syn group at 4 weeks. Lower limit of quantification (LLOQ) is specified in brackets after each cytokine. Data presented as mean±SD. α-Syn=rAAV-α-Syn+PFF. Control=rAAV-(-)+DPBS. ND=non-detected. Two-way ANOVA with Šídák multiple comparison test was used to compare strains (DA vs. DA.VRA4) and α-syn model (α-Syn vs. control).

| **CSF cytokines at 4 weeks (pg/ml)** | **DA Control (n=7)** | **DA.VRA4 Control (n=7)** | **DA α-Syn (n=7)** | **DA.VRA4 α-Syn (n=8)** |
|---|---|---|---|---|
| **IFNγ (39.7)** | 2.63±1.25 | 3.50±1.89 | 2.75±1.04 | 3.44±0.924 |
| **IL-10 (163)** | 4.67±1.74 | 3.71±1.23 | 5.65±2.40 | 5.03±2.29 |
| **IL-13 (12.5)** | ND | ND | ND | ND |
| **IL-1β (102)** | ND | ND | ND | 9.69±9.95 |
| **IL-4 (8.00)** | 0.415±0.268 | ND | ND | ND |
| **IL-5 (82.0)** | 23.1±4.88 | 21.8±4.30 | 23.4±9.15 | 24.0±8.82 |
| **IL-6 (96.9)** | 51.9±12.0 | 42.4±16.9 | 55.1±25.1 | 55.0±40.0 |
| **KC/GRO (21.7)** | 75.9±19.2 | 92.7±21.3 | 78.5±45.1 | 108±43.3 |
| **TNF (9.10)** | 0.954±0.326 | 1.21±0.338 | 1.13±0.550 | 2.21±2.24 |

**Table S3.** Cytokine levels in CSF from DA and DA.VRA4 rats in α-Syn or control groups at 8 weeks. Lower limit of quantification (LLOQ) is specified in brackets after each cytokine. Data presented as mean±SD. α-Syn=rAAV-α-Syn+PFF. Control=rAAV-(-)+DPBS. ND=non-detected. Two-way ANOVA with Šídák multiple comparison test was used to compare strains (DA vs. DA.VRA4) and α-syn model (α-Syn vs. control). ** $p < 0.01$ (DA vs. DA.VRA4). ## $p < 0.01$ and ### $p < 0.001$ (α-Syn vs. control).

| **CSF cytokines at 8 weeks (pg/ml)** | **DA Control (n=7)** | **DA.VRA4 Control (n=8)** | **DA α-Syn (n=8)** | **DA.VRA4 α-Syn (n=8)** |
|---|---|---|---|---|
| **IFNγ (39.7)** | ND | ND | 1.10±0.597 | 1.52±2.34 |
| **IL-10 (163)** | ND | ND | 1.75±0.738 | 3.16±1.07 ** |
| **IL-13 (12.5)** | ND | ND | ND | ND |
| **IL-1β (102)** | ND | ND | ND | ND |
| **IL-4 (8.00)** | ND | ND | 0.477±0.331 | 0.683±0.334 |
| **IL-5 (82.0)** | 9.60±4.75 | ND | 12.7±2.76 | 12.4±5.40 |
| **IL-6 (96.9)** | 12.0±12.0 | 6.56±7.65 | 44.6±20.8 ### | 31.0±11.2 ## |
| **KC/GRO (21.7)** | 50.0±25.1 | 79.9±67.1 | 60.1±30.1 | 51.8±6.37 |
| **TNF (9.10)** | 0.198±0.190 | 0.267±0.199 | 0.407±0.183 | 0.431±0.276 |

**Table S4.** Cytokine levels in serum from naïve DA and DA.VRA4 rats. Lower limit of quantification (LLOQ) is specified in brackets after each cytokine. Data presented as mean±SD. ND=non-detected. Unpaired Student's t-test. * $p < 0.05$.

| **Serum cytokines (pg/ml)** | **DA Naïve (n=6)** | **DA.VRA4 Naïve (n=6)** |
|---|---|---|
| **IFNγ (39.7)** | ND | ND |
| **IL-10 (163)** | 11.6±3.09 | 13.2±6.14 |
| **IL-13 (12.5)** | 1.10±1.09 | 1.15±0.695 |
| **IL-1β (102)** | 10.8±8.35 | 28.5±14.2 * |
| **IL-4 (8.00)** | 0.449±0.287 | ND |
| **IL-5 (82.0)** | 17.9±6.04 | 24.4±12.1 |
| **IL-6 (96.9)** | ND | 11.1±6.13 |
| **KC/GRO (21.7)** | 289±203 | 216±108 |
| **TNF (9.10)** | 3.34±0.564 | 4.22±0.567 * |

**Table S5.** Cytokine levels in serum from DA and DA.VRA4 rats in α-Syn and control groups at 4 weeks. Lower limit of quantification (LLOQ) is specified in brackets after each cytokine. Data presented as mean±SD. α-Syn=rAAV-α-Syn+PFF. Control=rAAV-(-)+DPBS. ND=non-detected. Two-way ANOVA with Šídák multiple comparison test was used to compare strains (DA vs. DA.VRA4) and α-syn model (α-Syn vs. control). * $p < 0.05$ and ** $p < 0.01$ (DA vs. DA.VRA4). # $p < 0.05$ (α-Syn vs. control).

| **Serum cytokines at 4 weeks (pg/ml)** | **DA Control (n=7)** | **DA.VRA4 Control (n=7)** | **DA α-Syn (n=7)** | **DA.VRA4 α-Syn (n=7)** |
|---|---|---|---|---|
| **IFNγ (39.7)** | ND | ND | ND | ND |
| **IL-10 (163)** | 14.8±4.07 | 10.6±3.37 | 15.9±4.98 | 13.8±3.77 |
| **IL-13 (12.5)** | ND | ND | ND | 1.13±1.07 |
| **IL-1β (102)** | ND | 14.4±5.81 | ND | 23.2±6.70 # |
| **IL-4 (8.00)** | 0.524±0.160 | 0.438±0.182 | 0.538±0.346 | 0.465±0.208 |
| **IL-5 (82.0)** | 25.1±6.85 | 28.7±4.18 ** | 24.8±10.8 | 37.3±5.78 |
| **IL-6 (96.9)** | 18.8±18.1 | ND | 20.4±10.4 | 20.9±16.2 |
| **KC/GRO (21.7)** | 334±312 | 240±69.3 | 163±48.2 | 221±62.4 |
| **TNF (9.10)** | 2.03±0.162 | 2.68±0.373 ** | 1.89±0.217 | 2.42±0.561 * |

**Table S6.** Cytokine levels in serum from DA and DA.VRA4 rats in α-Syn and control groups at 8 weeks. Lower limit of quantification (LLOQ) is specified in brackets after each cytokine. Data presented as mean±SD. α-Syn=rAAV-α-Syn+PFF. Control=rAAV-(-)+DPBS. ND=non-detected. Two-way ANOVA with Šídák multiple comparison test was used to compare strains (DA vs. DA.VRA4) and α-syn model (α-Syn vs. control). *** $p < 0.001$ (DA vs. DA.VRA4).

| **Serum cytokines at 8 weeks (pg/ml)** | **DA Control (n=7)** | **DA.VRA4 Control (n=7)** | **DA α-Syn (n=7)** | **DA.VRA4 α-Syn (n=7)** |
|---|---|---|---|---|
| **IFNγ (39.7)** | 7.33±12.0 | ND | 15.7±21.5 | 11.9±24.4 |
| **IL-10 (163)** | ND | ND | ND | ND |
| **IL-13 (12.5)** | 1.36±0.980 | 1.31±1.17 | 1.48±0.837 | 1.70±0.820 |
| **IL-1β (102)** | 36.8±30.2 | 44.2±19.5 | 24.1±16.0 | 47.0±27.8 |
| **IL-4 (8.00)** | ND | ND | ND | ND |
| **IL-5 (82.0)** | 26.7±19.0 | 38.1±31.6 | 24.8±12.1 | 35.2±20.0 |
| **IL-6 (96.9)** | 43.7±38.2 | 45.6±31.4 | 62.6±34.0 | 73.7±31.6 |
| **KC/GRO (21.7)** | 257±135 | 305±99.6 | 215±114 | 283±96.2 |
| **TNF (9.10)** | 3.12±0.362 | 4.33±0.584 *** | 3.10±0.432 | 4.30±0.393 *** |

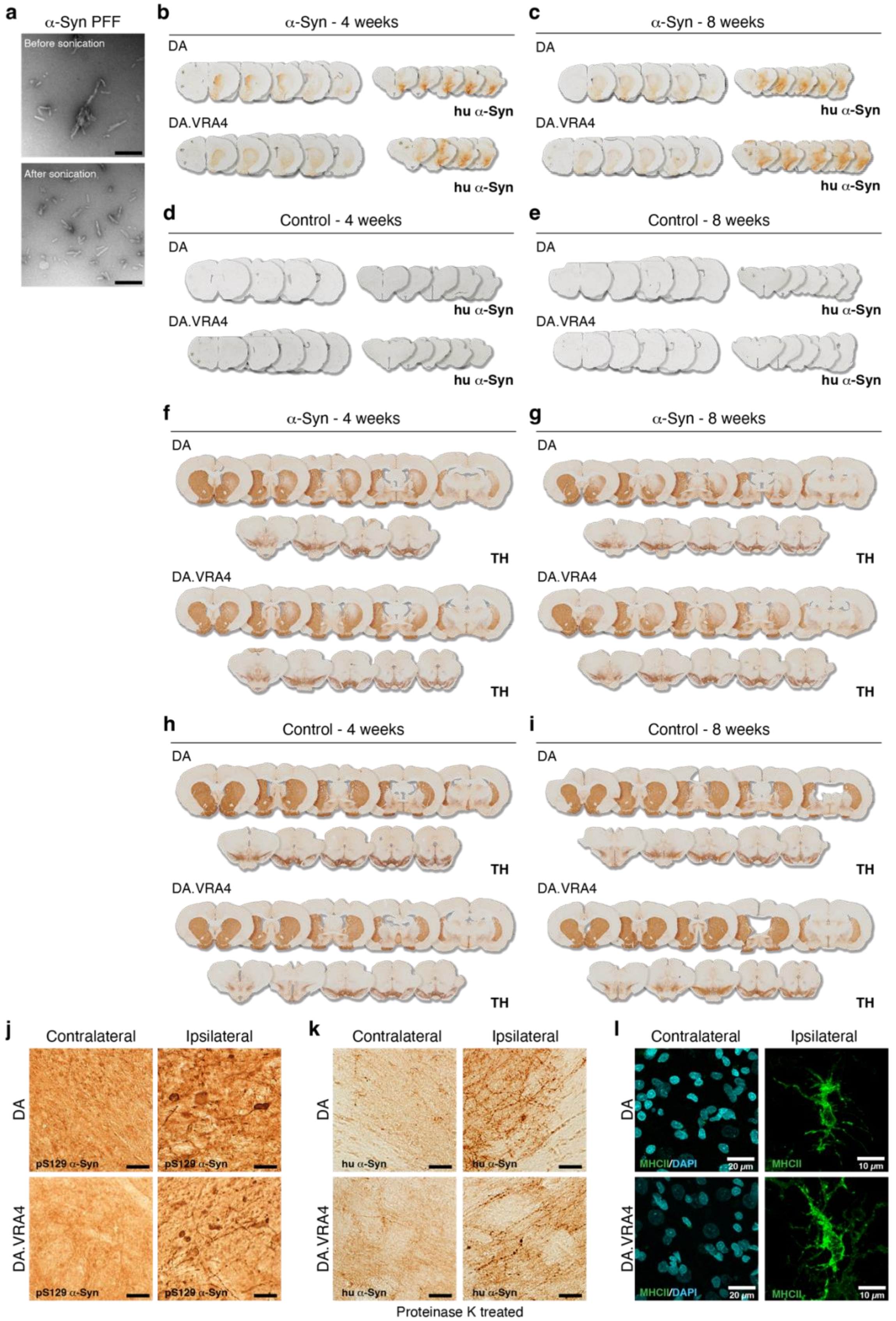


**Fig. S1 Representative images of human α-Syn and TH immunostaining in the brain of α-Syn and control groups. a.** TEM images of α-Syn PFF before (top) and after (bottom) sonication; sonicated PFF were used for striatal seeding. Scale bar = 200 nm. Expression of human α-Syn was detected at 4- (**b**) and 8-weeks (**c**) in the α-Syn groups. **d-e.** Control groups did not show any signal for human α-Syn at 4- or 8-weeks. **d.** Loss of TH signal was evident at 4- (**f**) and 8-weeks (**g**) in the α-Syn groups in both DA and DA.VRA4 rats. No TH loss was observed in control groups at 4- (**h**) or 8-weeks (**i**). Unilateral injection of rAAV-α-Syn+PFF resulted in pathological forms of α-Syn determined by pS129 α-Syn (**j**) and proteinase K resistant human α-Syn (**k**) stainings. **l**. Upregulation of MHCII on microglia was observed in the ipsilateral midbrain. Representative images in the α-Syn group at 8 weeks. **j-k.** Scale bar = 20 µm. **l**. Contralateral scale bar = 20 µm, ipsilateral scale bar = 10 µm. α-Syn=rAAV-α-Syn+PFF. Control=rAAV-(-)+DPBS.

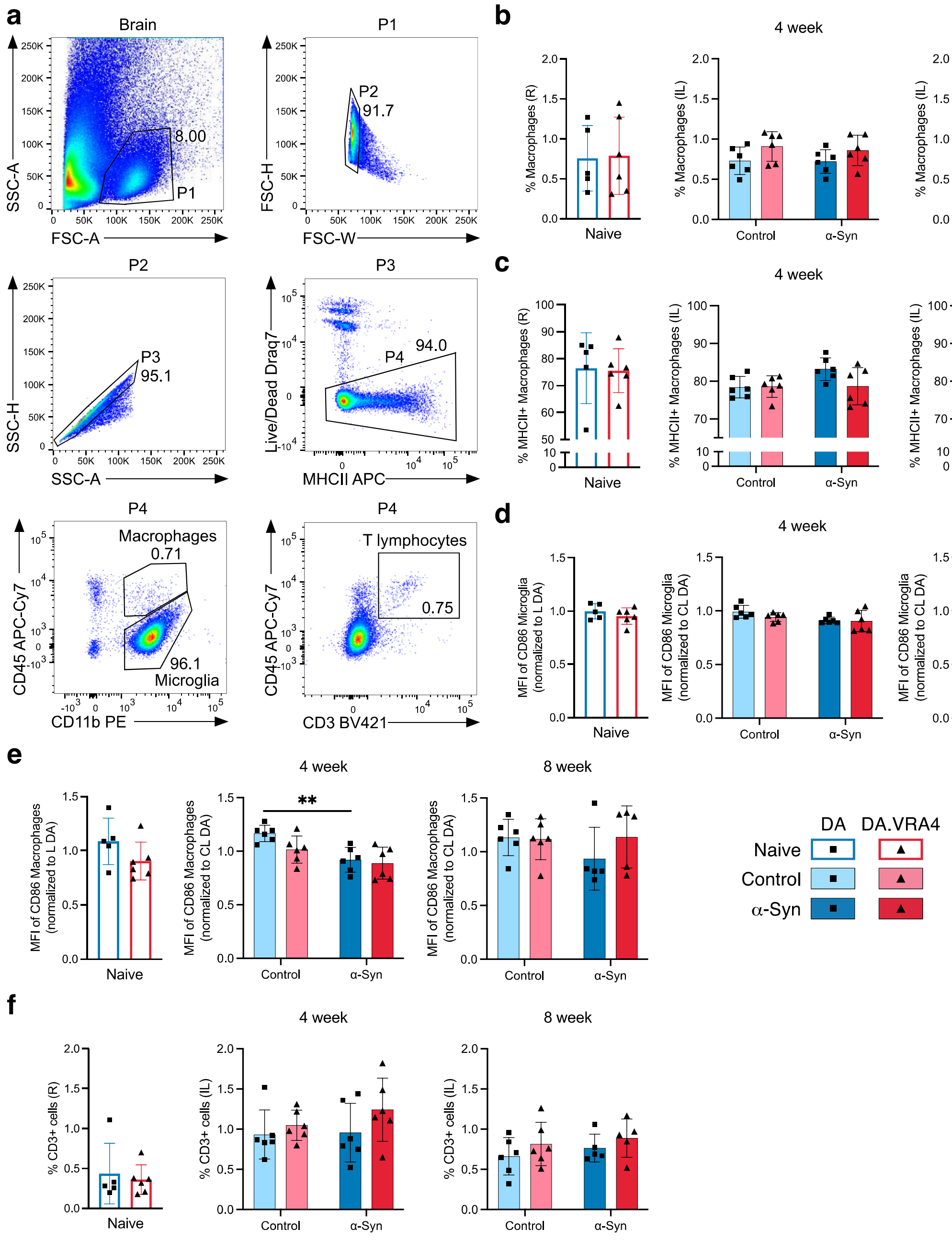


**Fig. S2 rAAV-α-Syn+PFF injection does not change brain-infiltrating macrophage/monocyte number, MHCII+ macrophage numbers or infiltration of lymphocytes. a.** Complete gating strategy of brain hemispheres for flow cytometry. **b.** Quantification of macrophages (CD45$^{high}$CD11b+) in right (R)/ipsilateral (IL) hemispheres. **c.** Quantification of MHCII+ microglia in the R/IL hemispheres. **d.** Quantification of relative median fluorescence intensity (MFI) of CD86 on microglia in R/IL hemispheres. At each recording session, R/IL MFI-values were normalized to the mean MFI-values in left(L)/contralateral (CL) hemispheres from DA rats **e.** MFI of CD86 on macrophages in R/IL hemispheres. **f.** CD3+ cells in IL hemisphere do not change in response to rAAV-α-Syn+PFF. Naïve (DA n=5, DA.VRA4 n=6), 4-week; control (DA n=6, DA.VRA4 n=6) and α-Syn (DA n=6, DA.VRA4 n=6), 8-week; control (DA n=6, DA.VRA4 n=6) and α-Syn (DA n=5, DA.VRA4 n=5). Data presented as mean ± SD with individual values. α-Syn=rAAV-α-Syn+PFF. Control=rAAV-(-)+DPBS. Naïve DA and DA.VRA4 rats were compared by unpaired Student's t-test. Two-way ANOVA with Šídák multiple comparison test was used to compare strains (DA vs. DA.VRA4) and experimental groups (α-Syn vs. control) at 4- and 8-weeks. $^{**}p < 0.01$

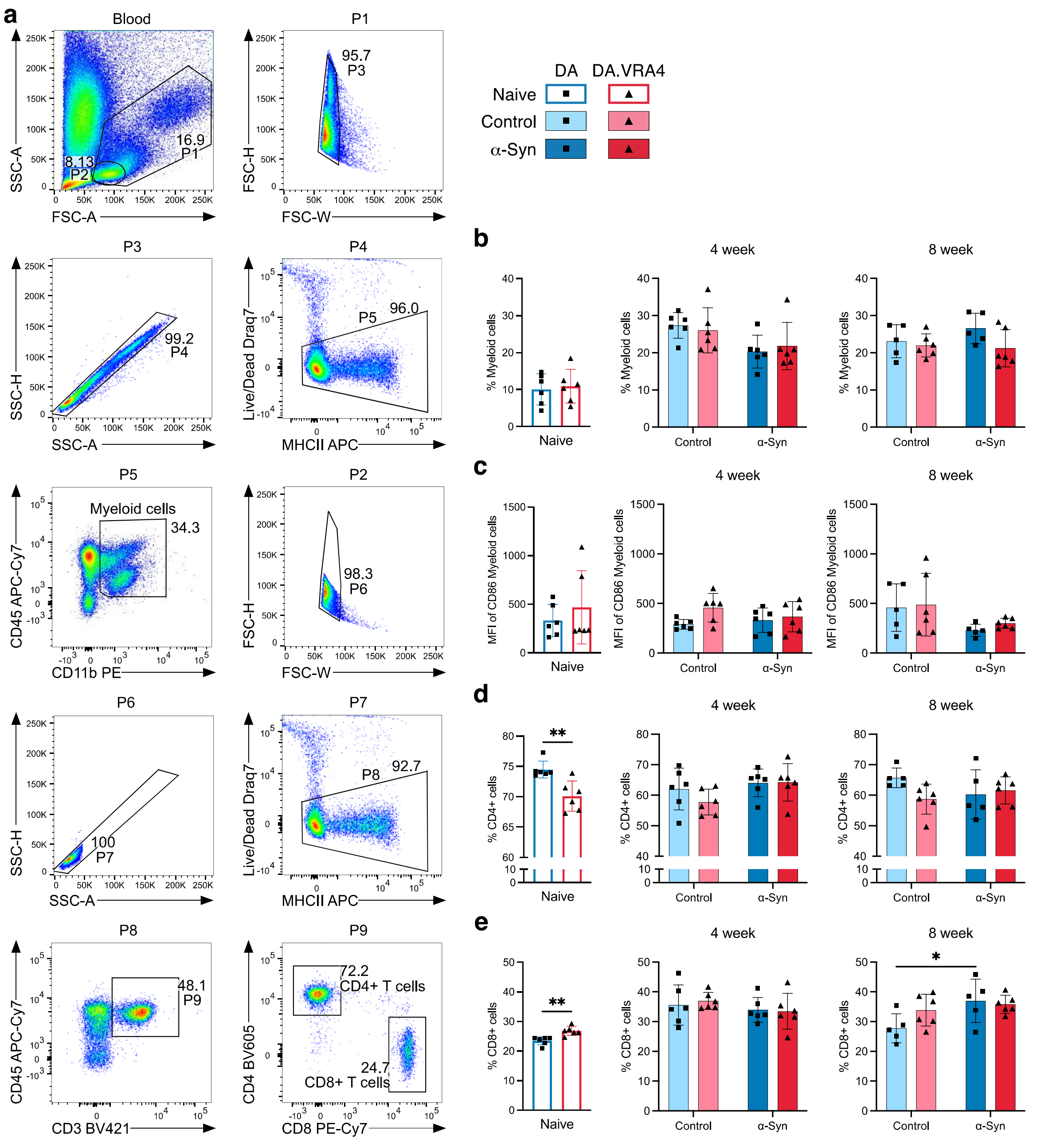


**Fig. S3 Circulating myeloid numbers and myeloid CD86 expression does not change in response to rAAV-α-Syn+PFF. a.** Gating strategy for blood flow cytometry. **b.** Percentage of myeloid cells (CD45+CD11b+) in blood. **c.** MFI of CD86 on myeloid cells. **d.** Percentage of CD4+ T lymphocytes. **e.** Percentage of CD8+ T lymphocytes. **b-e.** Naïve (DA n=6, DA.VRA4 n=6), 4 week; control (DA n=6, DA.VRA4 n=6) and α-Syn (DA n=6, DA.VRA4 n=6), 8 week; control (DA n=5, DA.VRA4 n=6) and α-Syn (DA n=5, DA.VRA4 n=6). Data presented as mean ± SD with individual values. α-Syn=rAAV-α-Syn+PFF. Control=rAAV-(-)+DPBS. Naïve DA and DA.VRA4 rats were compared by unpaired Student's t-test. Two-way ANOVA with Šídák multiple comparison test was used to compare strains (DA vs. DA.VRA4) and experimental groups (α-Syn vs. control) at 4- and 8-weeks. * $p < 0.05$ and ** $p < 0.01$.
